# SuperCell2.0 enables semi-supervised construction of multimodal metacell atlases

**DOI:** 10.64898/2026.02.19.706848

**Authors:** Léonard Hérault, Aurélie AG Gabriel, Benoît Duc, Bastien Dolfi, Aisha Shah, Johanna A. Joyce, David Gfeller

## Abstract

Multimodal single-cell atlases comprising hundreds of thousands of cells provide unique resources for exploring complex biological tissues and generating testable hypotheses. To streamline the analysis of such large datasets, we introduce SuperCell2.0, a robust workflow to build (semi-)supervised multimodal metacells. We demonstrate that multimodal metacells outperform metacells built with a single modality, improve inter-modality consistency, and facilitate integration of multiomic single-cell datasets. SuperCell2.0 can further leverage full or partial cell type annotations to improve metacell quality. This workflow enables us to construct multimodal metacell atlases from blood and tumor samples and identifies interferon-primed monocytes and macrophages in the circulation and in the tumor microenvironment. Markers derived from the metacell analysis enable us to sort and phenotypically characterize this population in healthy donors. Overall, our work demonstrates how SuperCell2.0 facilitates the analysis of large multimodal single-cell atlases.

## Introduction

Single-cell RNA sequencing (scRNA-seq) of hundreds of thousands of cells enables deep phenotypic characterization of complex tissues^1,2^. In cancer, large single-cell atlases have been generated across numerous tumor types^3–5^, providing detailed maps of the tumor microenvironment (TME), a dynamic ecosystem that shapes cancer progression and treatment response^6^. Recent technological advances now support richer molecular profiling through single-cell multiomics, in which multiple modalities are measured within the same cells^7,8^. For example, CITE-seq^9^ combines transcriptomes with surface protein measurements, and 10x Multiome^10^ jointly profiles chromatin accessibility and gene expression. Both technologies have been deployed in clinical studies, generating multimodal TME datasets containing tens of thousands of cells from dozens of patients^11–13^. These data hold promise for dissecting cell type–specific regulatory programs such as macrophage polarization^14^ and may ultimately improve our ability to therapeutically target the TME^15^.

Analysis of such large multimodal datasets requires robust computational methods capable of handling the considerable sparsity of these data, the high number of cells and samples, and the possible different origins of the samples^16,17^. This need is particularly acute for atlas-level studies, which integrate samples across multiple donors and anatomical sites and therefore exhibit substantial batch effects that must be corrected for reliable downstream analyses^18,19^.

To address these challenges, the concept of metacells, defined as disjoint and homogeneous groups of transcriptomically similar cells, was introduced for scRNA-seq20–24. Metacell-based strategies have since been successfully applied to large TME datasets^5,25–28^. The strategy consisting of an initial sample size reduction using metacells prior to batch correction has proven especially effective, improving both accuracy and computational efficiency^5,21,23,29^.

Recent work extended metacells to single-cell assay transposase accessible chromatin with sequencing (scATAC-seq) and cytometry data23,29–31. This paved the way to significant enhancements in single-cell multiomic data analyses. In particular, metacells have been widely used for gene regulatory network (GRN) analyses from single-cell multiomic data owing to the reduction in the dropout noise^32–37^.

Most existing metacell tools remain unimodal^20–24,30,31^, which is suboptimal for multimodal datasets where complementary modalities can resolve cell identity more accurately when analyzed jointly^29,38^. Moreover, metacell-based integration has rarely been evaluated beyond transcriptomic data. Considering all modalities is a valuable asset for large multimodal atlas analysis, where the different modalities may be affected by specific batch effects which need to be corrected, further increasing the computational load^38^.

Another limitation of current metacell tools is that they typically ignore prior cell-type annotations, grouping cells solely by molecular similarity. This can generate impure metacells that mix biologically distinct cell types^24^. In practice, obtaining complete and precise annotations is challenging, whereas coarse or partial annotations are now more accessible thanks to recent advances in automated cell annotation^39–41^. Akin to semi-supervised approaches used in other types of single-cell data analyses42–44, this prior knowledge could be used for the identification of metacells with increased purity.

Here we present SuperCell2.0, a semi-supervised metacell framework for single-cell (multi)modal data. SuperCell2.0 performs network-based coarse graining across modalities to merge highly similar cells into metacells suitable for quantitative multimodal analyses, including large-scale atlas integration. It can further leverage prior knowledge of partial cell annotation to improve metacell quality. Applying SuperCell2.0 to large CITE-seq and 10x Multiome atlases comprising hundreds of thousands of cells and dozens of donors enables us to characterize, across different modalities, interferon-primed macrophages and monocytes in the TME and in the peripheral blood. Building on this analysis, we derive reliable cell surface markers of interferon-primed monocytes that we experimentally validate in healthy donors.

## RESULTS

### SuperCell2.0 builds semi-supervised multimodal metacells

The unsupervised SuperCell2.0 workflow leverages the different modalities of a single-cell experiment to identify robust metacells (**Fig. 1A**). First, the dimension of each modality is reduced using approaches specific to each of these modalities. These include Principal Component Analysis (PCA) of highly variable genes for RNA, or proteins for antibody derived tags (ADTs), and Latent Semantic Indexing (LSI) of peaks for ATAC. Second, SuperCell2.0 takes as input these latent spaces to build a multimodal k-Nearest Neighbor (kNN) graph of the cells using the Weighted Nearest Neighbor (WNN) algorithm^38^. If only one modality is provided (e.g., RNA), SuperCell2.0 builds a standard kNN from the corresponding latent space (e.g., PCA). Third, SuperCell2.0 identifies metacells on the kNN graph using random walks weighted by cell-to-cell affinities with the walktrap algorithm^45^. Fourth, single cells are grouped into metacells at a user-defined graining level γ, which corresponds to the ratio of the number of single cells to the number of desired metacells. Single-cell data are aggregated by summing raw counts in each metacell for each modality.

**Figure 1:**
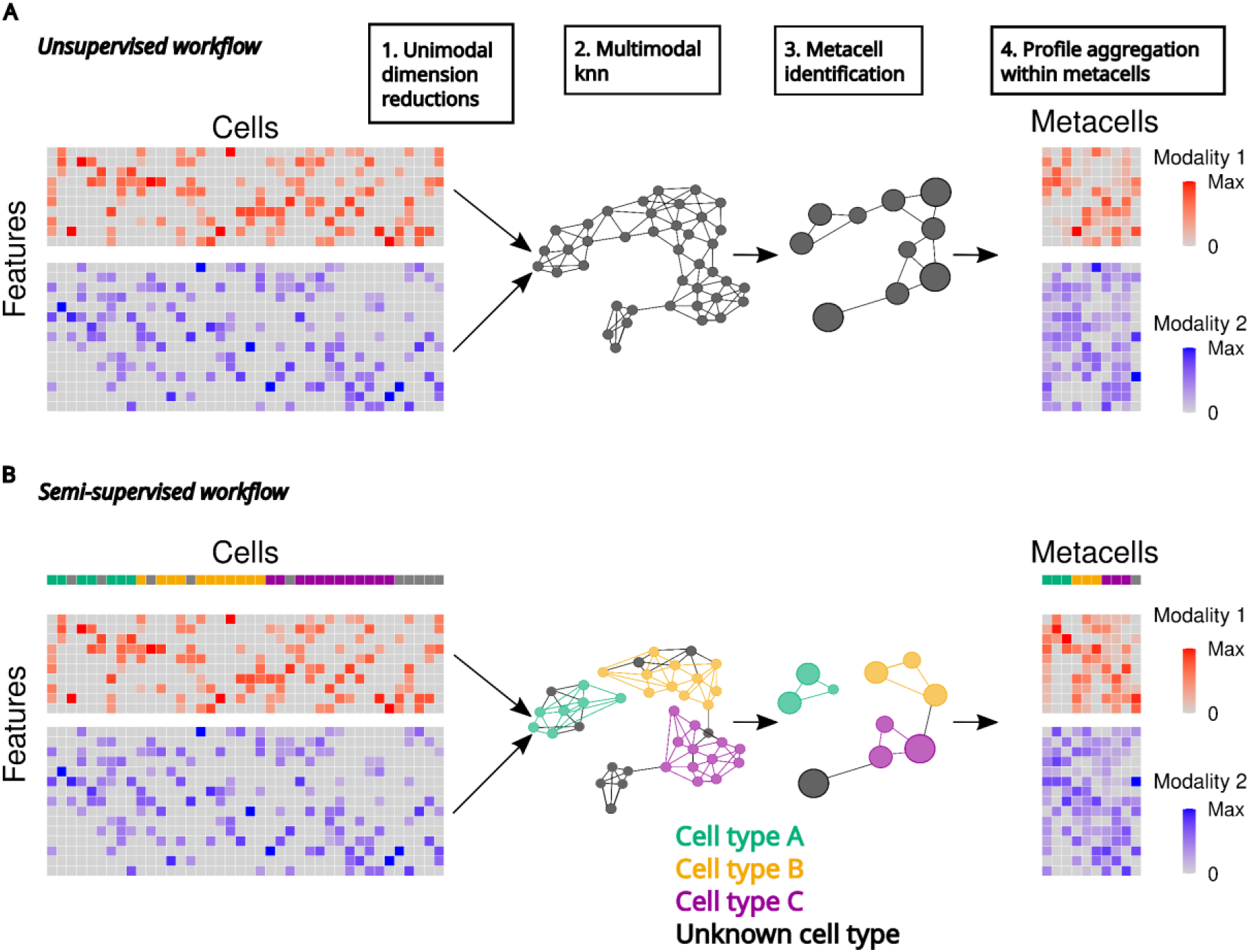
SuperCell2.0 builds semi-supervised multimodal metacells. **A** Unsupervised workflow of SuperCell2.0 for single-cell multimodal data with two modalities. **B** Semi-supervised workflow of SuperCell2.0 for single-cell multimodal data with cells partially annotated to three different cell types.

SuperCell2.0 can also leverage complete or partial single-cell annotation to guide metacell identification and limit metacell impurities. To achieve this, the semi-supervised workflow of SuperCell2.0 consists of using single-cell annotation to build kNN graphs for each annotated cell type separately. Then, unannotated cells are connected to these graphs using edges of unannotated cells from the unsupervised kNN of all cells (**Fig. 1B**). The resulting graph is used as input for the metacell identification with the walktrap algorithm. The annotation used in the (semi-)supervised workflow can come from automated single-cell annotation tools or previous single-cell manual annotation.

### SuperCell2.0 identifies robust multimodal metacells

We first applied the unsupervised SuperCell2.0 workflow to a Peripheral Blood Mononuclear Cell (PBMC) 10x Multiome dataset comprising joint RNA and ATAC measurements for 10,412 single nuclei (**Table 1**). Using a graining level of γ = 75, we obtained 138 metacells that homogeneously covered the single-cell space (**Fig. 2A and Fig. S1A**). Our metacells substantially reduced dropout noise in both modalities, resulting in a higher number of detected genes and peaks compared with single cells (**Fig. 2B**).

**Figure 2:**
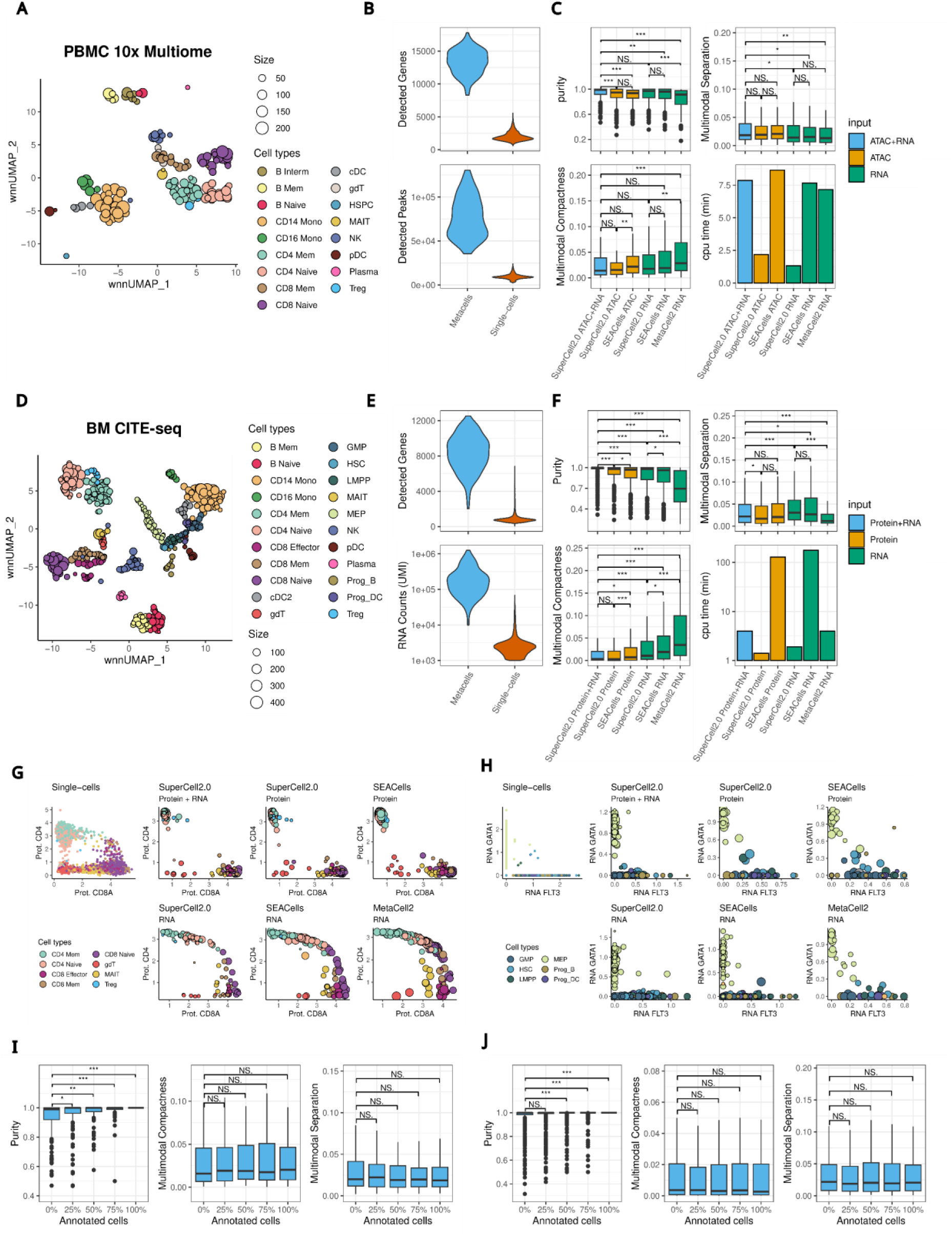
SuperCell2.0 identifies robust multimodal metacells and leverages partial prior knowledge to increase metacell quality. **A** Coverage plots of metacells identified by the unsupervised multimodal SuperCell2.0 workflow on the PBMC 10x Multiome dataset at γ=75. **B** Distribution of the numbers of detected genes and peaks per metacell and single cell (PBMC 10x Multiome dataset). **C** Metacell purity, multimodal separation and compactness and CPU time of SuperCell2.0 (unsupervised), SEACells and MetaCell2 using as input ATAC+RNA, ATAC only and RNA only for the PBMC 10x Multiome data. For clarity, outliers are shown only for purity. **D** Coverage plots of metacells identified by the unsupervised multimodal SuperCell2.0 workflow on BM CITE-seq data at γ=75. **E** Distribution of the number of detected genes and RNA (UMI) counts per metacell and single cell (BM CITE-seq dataset). **F** Metacell purity, multimodal separation and compactness (outliers not shown), CPU time of SuperCell2.0 (unsupervised), SEACells and MetaCell2 using as input Protein with RNA, Protein only and RNA only. **G** CD8A and CD4 protein expression in T lymphocyte metacells identified with SuperCell2.0, SEACells and MetaCell2 using as input Protein & RNA, Protein only, and RNA only from the BM CITE-seq dataset. H *FLT3* and *GATA1* RNA expression in progenitor metacells identified with SuperCell2.0, SEACells and MetaCell2 using as input Protein & RNA, Protein only and RNA only from the BM CITE-seq dataset. **I** Purity, multimodal compactness and separation of metacells identified at different percentages of annotated cells on the PBMC 10x Multiome data with the semi-supervised workflow of SuperCell2.0. **J** Purity, multimodal compactness and separation of metacells identified at different percentages of annotated cells on the BM CITE-seq data with the semi-supervised and multimodal workflow of SuperCell2.0. Differences in metric distributions were tested using Wilcoxon tests in **C, F, I** and **J** with results grouped into (***) for p-values < 0.001, (**) for p-values < 0.01, (*) for p-values < 0.05 and no significant difference (NS) for p-value > 0.05.

**Table 1:**
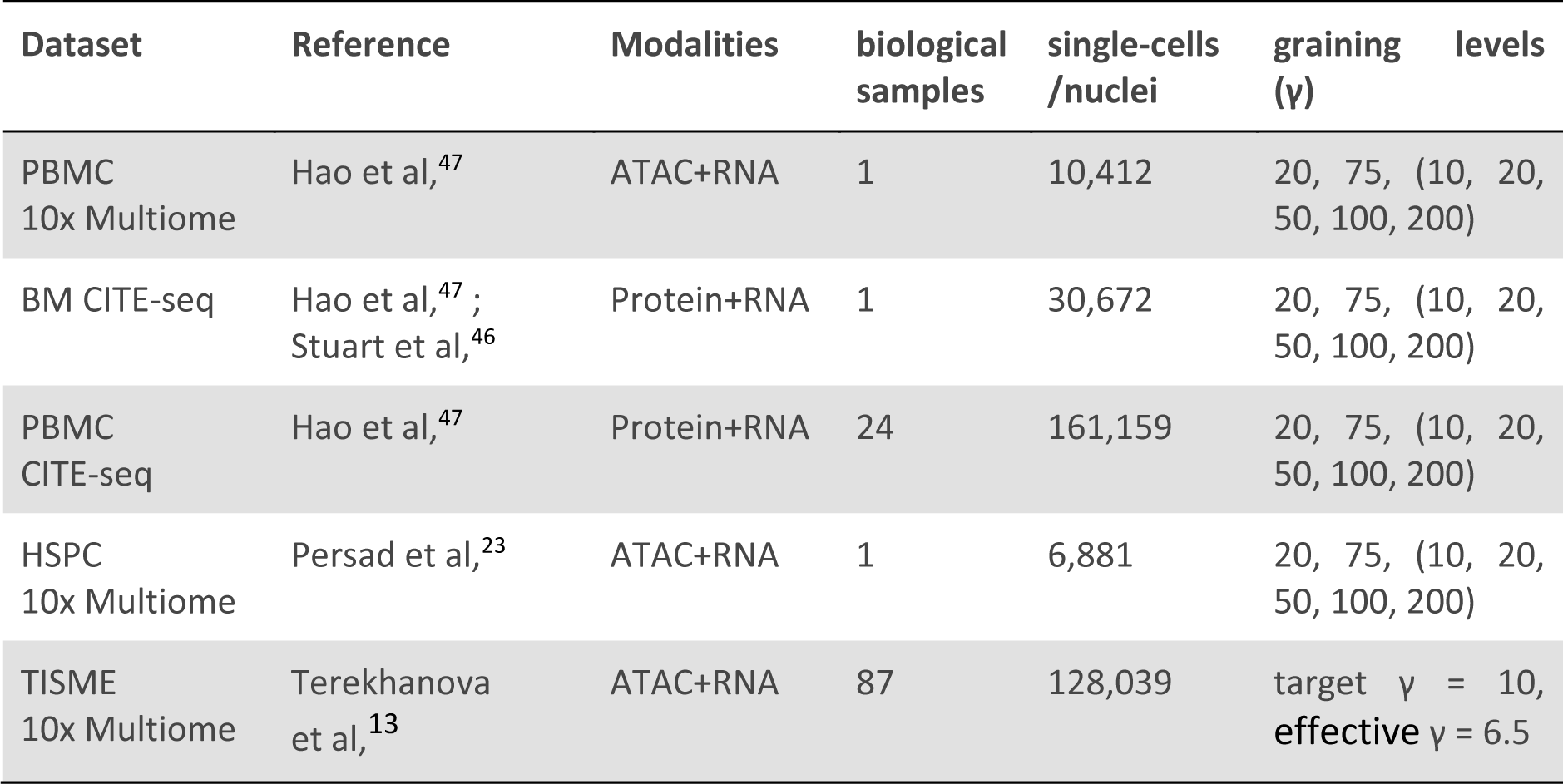
Summary of the datasets used in this study. Graining levels in parentheses were used only for the analysis of intermodality correlation/GRN inference.

To benchmark the unsupervised multimodal SuperCell2.0 workflow, we compared it with SEACells23, which identifies metacells on either RNA or ATAC alone, and MetaCell222, which relies solely on RNA. For completeness, we also included SuperCell2.0 applied to RNA alone or ATAC alone. All methods were evaluated at γ = 75 using established metrics of purity, compactness, and separation^21,23,24^ (see Methods). Multimodal compactness (lower is better) and separation (higher is better) were computed as the average of scaled RNA and ATAC metrics.

On average, metacells generated by the multimodal SuperCell2.0 workflow were purer and provided better coverage of the single-cell space than those obtained from any unimodal approach (**Fig. 2C and Fig. S1A**). Multimodal metacells were also more compact than those produced by MetaCell2, while showing comparable compactness to other approaches (**Fig. 2C and Fig. S1B**). Moreover, the multimodal workflow yielded better-separated metacells than all RNA-only methods (**Fig. 2C and Fig. S1B**). Metacell size distributions were similar between SuperCell2.0 and SEACells (**Fig. S1C**).

Regarding computational efficiency, the SuperCell2.0 unimodal workflow was faster than SEACells and MetaCell2 while the SuperCell2.0 multimodal workflow performed on par with other methods applied to single modalities (**Fig. 2C**). MetaCell2, which requires no preprocessing, used the least memory (**Fig. S1D**). Repeating the benchmark at γ = 20 produced consistent results for both metacell quality and computational efficiency, with the exception that SEACells required approximately six times longer CPU time than the other approaches at this lower γ (**Fig. S1E–J**).

We next applied SuperCell2.0 to a bone marrow (BM) CITE-seq dataset comprising 30,672 progenitor and mature immune cells^46^ (**Table 1**). At a graining level of γ = 75, we obtained 408 metacells that showed excellent coverage of the multimodal single-cell space (**Fig. 2D** and **Fig. S2A**). Metacell substantially reduced RNA dropout noise, yielding higher unique molecular identifier (UMI) counts and more detected genes compared to single cells (**Fig. 2E**).

As before, we benchmarked the multimodal SuperCell2.0 workflow (Protein + RNA) against unimodal approaches: SuperCell2.0 and SEACells using either Protein or RNA alone, and MetaCell2 using RNA. The multimodal workflow produced significantly purer metacells than all unimodal methods (**Fig. 2F**). It also generated, together with SuperCell2.0 using Protein alone, more compact metacells than the remaining approaches (**Fig. 2F and Fig. S2B**). RNA-based metacell identification with SuperCell2.0 and SEACells yielded the best-separated metacells but showed poor coverage of the single-cell space, consistent with their lower purity relative to the multimodal and protein-based approaches (**Fig. 2F and Fig. S2A–C**). In terms of computational performance, SuperCell2.0 was faster than SEACells and more memory-efficient than MetaCell2 and SEACells (**Fig. 2F and Fig. S2D**). Benchmarking at γ = 20 produced similar results, with SuperCell2.0 again achieving superior metacell quality and computational efficiency (**Fig. S2E–J**).

To further investigate differences in metacell purity, we examined the protein expression of CD8A and CD4 in metacells obtained with each method at γ = 75. The multimodal SuperCell2.0 workflow, as well as the protein-based unimodal implementations of SuperCell2.0 and SEACells, correctly segregated these markers into distinct CD4 and CD8 metacells. In contrast, RNA-based unimodal approaches produced numerous double-positive CD4/CD8A metacells with low purity, reflecting a mixture of the two cell types (**Fig. 2G and Fig. S2K**).

We next assessed RNA expression of *GATA1*, a marker of megakaryocytic and erythroid progenitors (MEPs), and *FLT3*, a marker of lymphoid and myeloid progenitors—including lympho-myeloid primed progenitors (LMPPs), B-cell and dendritic cell (DC) progenitors, and granulocyte–monocyte progenitors (GMPs). The multimodal SuperCell2.0 workflow and the RNA-based unimodal approaches SuperCell2.0 and SEACells correctly segregated these progenitor populations (**Fig. 2H and Fig. S2K**). Protein-based unimodal approaches and MetaCell2 failed to do so.

Overall, unimodal approaches were suboptimal for this CITE-seq dataset, in which mature immune cell identity is better captured by protein measurements, whereas progenitor states are more strongly defined by RNA expression, a pattern reflected in the single-cell WNN modality weights (**Fig. S2L**).

Altogether, our benchmark demonstrates that the multimodal unsupervised SuperCell2.0 workflow identifies robust metacells from the two most widely used single-cell multimodal platforms and outperforms common unimodal metacell construction tools. When restricted to a single modality, SuperCell2.0 performs significantly better than MetaCell2 and comparably to SEACells, while requiring fewer computational resources.

### SuperCell2.0 can integrate partial prior knowledge in multimodal metacell construction

Next, we assessed to what extent SuperCell2.0 can leverage partial single-cell annotations to improve metacell quality. We applied the multimodal semi-supervised workflow at γ = 75 while varying the proportion of annotated cells. As expected, metacell purity increased markedly as more annotations were provided, whereas compactness and separation remained largely unchanged for both the PBMC 10x Multiome (**Fig. 2I and Fig. S3A**) and BM CITE-seq datasets (**Fig. 2J and Fig. S3B**), indicating that the added prior knowledge complements rather than replaces the information captured by the unsupervised workflow.

Because unsupervised metacells were already highly pure in the BM CITE-seq dataset, we also tested the semi-supervised approach using RNA alone (**Fig. S3C**). In this setting, we observed significant gains in both purity and compactness, accompanied by a modest reduction in separation (**Fig. S3D**). With 75% of cells annotated, the approach clearly resolved CD4 and CD8 populations (**Fig. S3E**).

Overall, these results demonstrate that SuperCell2.0 can effectively leverage partial annotations to further improve metacell quality, primarily by increasing purity while preserving compactness and separation. The benefits of semi-supervision are particularly evident in settings with limited modality information or lower initial purity.

### Multimodal metacells improve inter-modality consistency

We next investigated whether multimodal metacells improve intermodality consistency in single-cell multimodal data. We first examined intermodality correlations in metacells obtained from the unsupervised multimodal SuperCell2.0 workflow at γ = 75 from the BM CITE-seq dataset. We observed stronger correlations between RNA and protein expression for CD8A and CD14 than in single cells, where RNA measurements are substantially affected by dropout (**Fig. 3A**). We next analyzed the entire antibody panel while varying γ from 10 to 200, using down-sampled single cells and randomly generated metacells as controls. Across most of the 25 measured proteins, RNA–protein correlations increased in metacells, demonstrating a clear gain in intermodality consistency (**Fig. 3B**).

**Figure 3:**
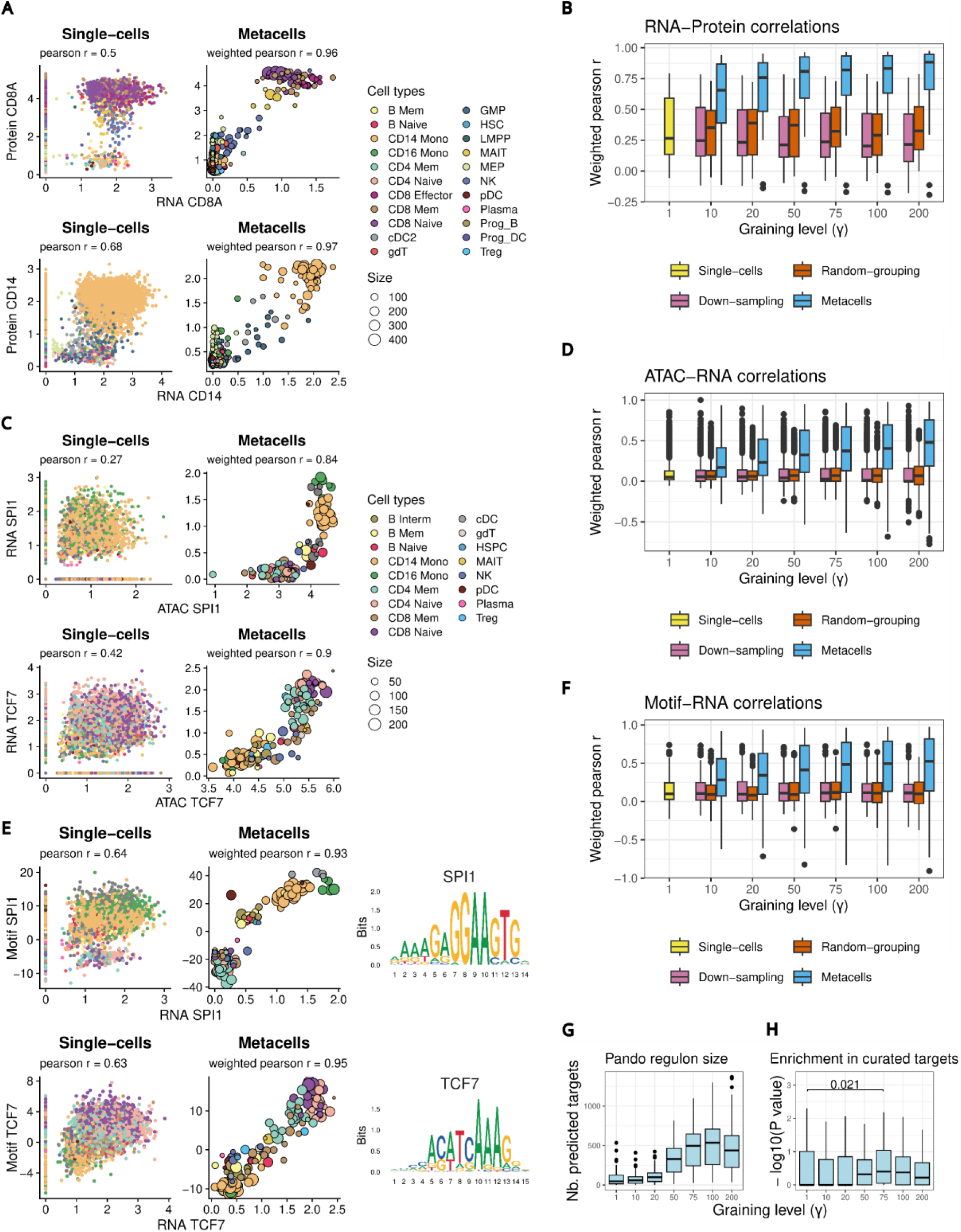
Multimodal metacells improve intermodality consistency. **A** RNA and surface protein expression in single cells and unsupervised multimodal metacells (γ=75) for CD8A and CD14 in the BM CITE-seq data. **B** Distribution of weighted Pearson RNA-Protein correlations at increasing γ for the 25 measured proteins in the BM CITE-seq data. Down-sampling and random grouping of single cells were performed as control for each γ. **C** Gene activity (ATAC) and RNA expression correlation in single cells and metacell (γ=75) for *SPI1* and *TCF7* in the PBMC 10x Multiome data. **D** Distribution of weighted Pearson gene activity - gene expression (ATAC-RNA) correlations at increasing γ for 4000 highly variable genes at the single-cell RNA level in the PBMC 10x Multiome data. Down-sampling (subsetting) and random grouping of single cells were performed as control for each γ. **E** TF motif accessibility (chromVAR) and TF RNA expression correlation in single cells and metacell for *SPI1* and *TCF7* in the PBMC 10x Multiome data. Corresponding TF motifs are represented as logos (right). **F** Distribution of weighted Pearson TF motif accessibility - gene expression correlations at different γ for TF markers of a PBMC cell type at the single-cell level. Down-sampling and random grouping of single cells were performed as controls for each γ. **G** Sizes of regulons (i.e. number of targets) inferred with Pando at increasing γ. **H** Pando regulon enrichments (-log10 P-value hypergeometric test) in curated targets from the CollecTRI database at increasing γ. Outliers are not shown. Differences in enrichment between γ from 10 to 200 (i.e., metacells) and γ=1 (i.e., single cells) were tested using Wilcoxon tests. (*) for p values < 0.05.

We then analyzed a human sample of a PBMC CITE-seq atlas^47^ of approximately 5,000 cells with a panel of 200 antibodies to assess reproducibility across biological contexts and a broader feature space (**Fig. S4A and Table 1**). Again, we observed enhanced RNA–protein consistency for CD8A and CD14, as well as globally across the entire protein panel (**Fig. S4B&C**).

We next assessed intermodality consistency in PBMC 10x Multiome data by examining the correlation between ATAC-derived gene body accessibility (gene activity) and RNA expression. Metacells showed higher correlations for *SPI1* (a myeloid transcription factor) and *TCF7* (a naïve T-cell transcription factor) than single cells, in which both modalities suffer from dropout noise (**Fig. 3C**). When extended to the top 4,000 highly variable genes for which gene activity scores could be computed, the analysis showed increasing ATAC–RNA correlations with higher γ, demonstrating improved intermodality consistency at the metacell level (**Fig. 3D**). These results are consistent with previous observations obtained with metacells built on a single modality^23^.

We obtained similar results on a Human Stem and Progenitor Cells (HSPC) 10x Multiome dataset^23^ analyzed with SuperCell2.0 (**Fig. S4D-F and Table 1**). Correlations between ATAC and RNA were generally lower in this dataset than in PBMCs (**Fig. S4G**). This may reflect the well characterized delay for gene opening/transcription in the differentiating HSPCs compared to PBMCs that are in a more stable state^48^.

To assess intermodality consistency at the regulatory level, we examined correlations between transcription factor (TF) motif accessibility in ATAC and corresponding TF RNA expression in the PBMC 10x Multiome dataset. We observed an increase in correlations for *SPI1* and *TCF7* (**Fig. 3E**). Considering a broader set of TFs identified as cell-type markers for both RNA expression and motif accessibility, we obtained progressively stronger correlations as γ increased (**Fig. 3F**).

To advance beyond paired feature correlations across modalities, we examined whether GRN inference from single-cell multiomic data benefits from the improved intermodality consistency provided by metacells. To this end, we applied the Pando approach^33^ to the PBMC 10x Multiome dataset across different γ. Regulon (i.e. inferred modules of a TF and its targets) size increased with γ, plateauing around γ = 100 (**Fig. 3G**). In parallel, regulons showed increased enrichment for curated TF targets from the CollecTRI database^49^ starting at γ = 50, with a peak at γ = 75 where the improvement relative to single cells was significant (**Fig. 3H**).

Together, these results demonstrate that metacells substantially improve intermodality consistency in both CITE-seq (RNA–protein) and 10x Multiome (ATAC–RNA) datasets across three distinct biological systems (BM, PBMC, and HSPC).

### Semi-supervised multimodal metacells enable integration of large multimodal cell atlases

Building and analyzing large multimodal cell atlases comprising samples from diverse donors and sites remains challenging due to the size of the data and substantial batch effects across modalities. Here, we developed and evaluated a (semi-)supervised workflow that leverages metacells and prior knowledge to address these challenges, with a focus on scalability and accurate batch effect correction. Our integration workflow begins with metacell identification from multimodal data in each atlas sample using SuperCell2.0, followed by modality-specific batch correction with the anchor-based method STACAS^42^ (**Fig. 4A**). Both steps can be (semi-)supervised using partial cell type annotations. Finally, the batch-corrected modalities are integrated using the WNN approach at the metacell level (**Fig. 4A**).

**Figure 4:**
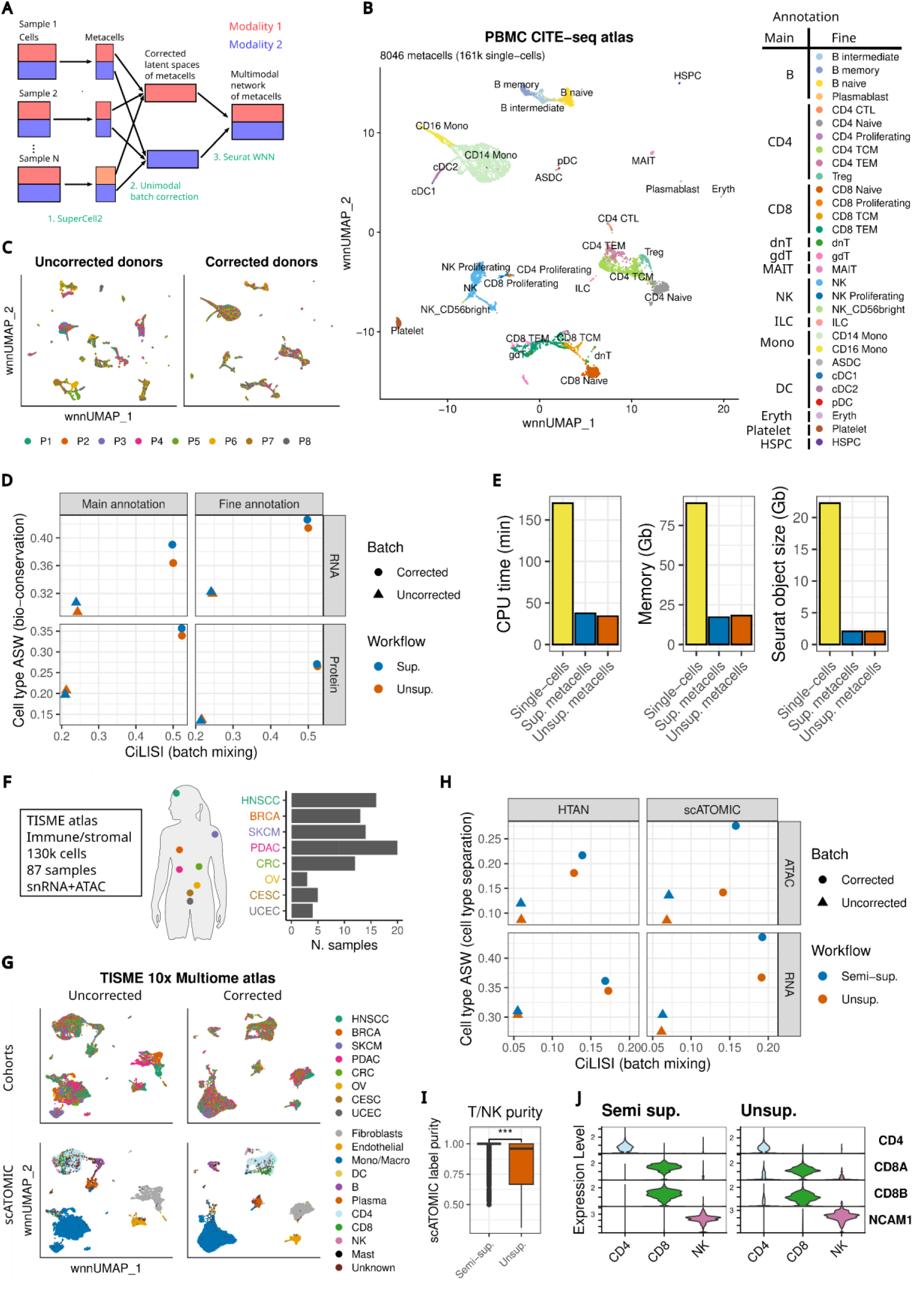
(Semi-)supervised multimodal metacells enable integration of large multimodal atlases. **A** Workflow to build large multimodal metacell atlases. Metacell identification (step 1) and unimodal batch correction (step 2) can be guided using (partial) cell annotations. **B** WNN UMAP of the integrated PBMC CITE-seq metacell atlas using the supervised workflow. Metacells are colored with the fine annotation (aggregated single-cell labels). **C** WNN UMAPs of the PBMC CITE-seq atlas, before and after batch correction of RNA and protein latent space using the supervised workflow. Metacells are colored by donors. **D** Cell type ASW (cell type separation) and CiLISI (batch mixing) of integrated metacells using the supervised or the unsupervised workflow to build the PBMC CITE-seq atlas. The metrics are computed using main and fine annotations in RNA and Protein latent spaces before and after batch correction. **E** Computational load of the metacell supervised workflow compared to the original single-cell workflow (anchor-based batch correction using Seurat). **F** 10x Multiome data (snRNA+ATAC) from HTAN. **G** WNN UMAPs of the TISME 10x Multiome atlas before and after batch correction of RNA and ATAC latent spaces using the supervised workflow. Metacells are colored by cancer type cohort and scATOMIC annotation (aggregated single-cell labels). **H** Cell type ASW (cell type separation) and CiLISI (batch mixing) of integrated metacells using the semi-supervised or the unsupervised workflow to build the TISME 10x Multiome atlas. The metrics were computed using scATOMIC and original HTAN labels in RNA and ATAC latent spaces before and after batch correction. **I** T/NK metacell purities in scATOMIC labels. Metacells were obtained with the semi-supervised and the unsupervised workflow. The purity difference was tested using a Wilcoxon test, (***) indicates a p-value < 0.001. **J** Violin plot of T/NK metacell expression of key marker genes obtained with the supervised and the unsupervised workflow.

We first assessed how effectively our workflow leverages prior knowledge when integrating a PBMC CITE-seq atlas of 24 samples comprising 161,159 cells annotated at both main and fine cell type levels^47^ (**Table 1**). The main cell type annotation was used to supervise the identification and integration of 8,046 metacells (γ = 20). After integration, we observed clear clustering of fine and main cell types at the metacell level (**Fig. 4B & Fig. S5A**), accompanied by improved batch mixing (**Fig. 4C & Fig. S5A**), indicating successful integration. Supervision with main cell type labels increased metacell purity relative to a fully unsupervised workflow (**Fig. S5B**), which also showed reduced batch effects (**Fig. S5C&D**). We next quantified batch correction at the two annotation levels in RNA and protein latent spaces using the cell type Average Silhouette Width (ASW) for cell type separation and the Cell-type integration Local Inverse Simpson’s Index (CiLISI) for batch mixing. For both modalities and both annotation levels, the supervised workflow achieved improved batch correction (higher ASW and CiLISI) than the unsupervised workflow (**Fig. 4D**). Regarding computational efficiency, we observed a substantial reduction in runtime and memory usage compared to the single-cell workflow (**Fig. 4E**). Integration of 160,000 multimodal single cells into 8,000 metacells can be performed on a laptop with less than 20 GB RAM, and the resulting R object is reduced 10-fold in size (20 GB to 2 GB) compared to single cells, thereby greatly facilitating downstream analyses **(Fig. 4E**). When benchmarking the workflow under partial annotation, the semi-supervised mode maintained performance close to the fully supervised mode with up to 80% annotated cells, both for metacell purity and batch effect correction (**Fig. S5E&F**).

We next evaluated our workflow on a single-cell multiomics dataset with limited prior knowledge: the 10x Multiome (RNA + ATAC in single nuclei) data from the Human Tumor Atlas Network (HTAN)13,50,51. Using scATOMIC41, we annotated and collected 130,000 immune and stromal cells from 87 tumor samples across 79 donors and 8 cancer types to construct a pan-cancer multiomic metacell atlas of the tumor immune and stromal microenvironment (TISME) (**Fig. 4F and Table 1**). Most cells (95%) could be assigned to immune and stromal subtypes, and this computational annotation showed strong concordance with the original coarse HTAN annotation, which we retained for external validation (**Fig. S6A**). We then relied on the scATOMIC annotation to guide the semi-supervised workflow. Using a target γ = 10 and a minimum of 200 metacells per sample, the semi-supervised SuperCell2.0 workflow yielded roughly 20,000 metacells (effective γ = 6.5). Multimodal WNN UMAPs before and after integration demonstrated successful batch correction, with improved cell type separation and batch mixing (including increased mixing across cancer cohorts) for both semi-supervised (**Fig. 4G & Fig. S6B**) and unsupervised workflows (**Fig. S6C**). Across RNA and ATAC modalities and for both the predefined HTAN and the predicted scATOMIC annotations, our benchmark showed that the best batch correction (highest cell type ASW and high CiLISI) was achieved with the semi-supervised workflow (**Fig. 4H**). Metacell purity was similar for the HTAN annotation, whereas scATOMIC-based purity was slightly higher in semi-supervised metacells **(Fig. S6D**). At the level of individual cell types, the semi-supervised workflow produced higher-purity metacells for T/NK subsets subdivided into CD4 T, CD8 T, and NK cells by scATOMIC (all grouped in the HTAN annotation), compared to the unsupervised metacells (**Fig. 4I & Fig. S6E**). This was supported by lower residual expression of *CD4*, *CD8A*, *CD8B*, and *NCAM1* in cell types where these markers are not expected **(Fig. 4J**). We also noted a significant reduction in computational load relative to single-cell integration (**Fig. S6F**).

In summary, these two applications show that our (semi-)supervised metacell workflow efficiently corrects batch effects across modalities and effectively leverages partial cell type annotations to build large multimodal metacell atlases.

### Multimodal metacell analysis of the TISME atlas reveals properties of interferon-responding macrophages

To illustrate how multiomic atlases can be used for data exploration at the metacell level, we performed a multimodal clustering of the TISME 10x Multiome atlas (see Methods). This yielded 40 metacell clusters that we grouped into ten major immune and stromal types (**Fig. 5A, S7A–D**). Stromal clusters consisted of endothelial, fibroblast, and mural cell types, whereas immune clusters comprised DCs, monocytes/macrophages, plasma cells, B cells, T cells, NK cells, and mast cells. These major types were defined by both RNA expression and chromatin accessibility around the transcriptional start sites (TSS) of representative marker genes (**Fig. 5B, Table S1-2**), and were found in different proportions across cohorts (**Fig. 5C**).

**Figure 5:**
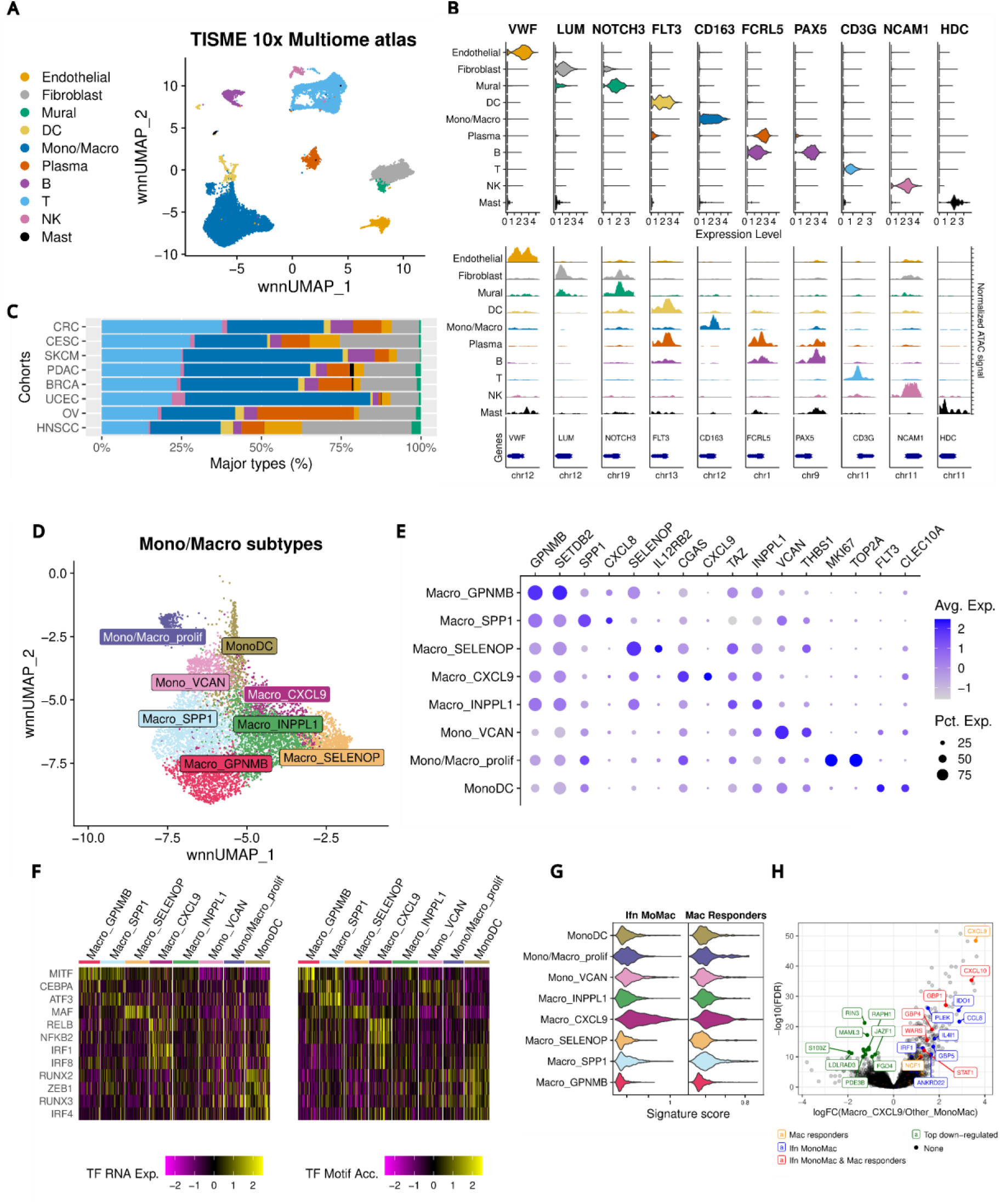
Multimodal metacell analysis of a TISME 10x Multiome atlas reveals properties of interferon-responding macrophages. **A** WNN UMAP of the TISME 10x Multiome atlas with metacells colored by major cell types. **B** Markers of major cell types at the RNA level with violin plots (upper panel) and at the ATAC level with coverage plots around the TSS (+/- 500 bp, lower panel). **C** Major cell type composition of the different cohorts of cancer types. **D** WNN UMAP of the tumor associated monocytes and macrophages subtypes. **E** Dot plots of RNA markers of monocytes and macrophages subtypes. **F** Pseudobulk heatmap of top TFs markers of monocytes and macrophages subtypes at the RNA and motif accessibility (ATAC) level. Pseudobulk data correspond to averaged RNA expression and motif accessibility in each subtype and each sample. **G** Score of signatures of Interferon response and ACT-TILs response in monocytes/macrophages. **H** Volcano plots of DEGs in CXCL9-high macrophages versus other tumor-associated monocytes/macrophages. Top genes of the signatures are highlighted.

We then annotated each cluster to a fine subtype using previously reported markers (**Fig. S7E**, see Methods). In particular, we achieved high-resolution classification of the monocyte/macrophage compartment (**Fig. 5D–E**, **Table S3-4**), as well as the T/NK and DC compartments **(Fig. S8A–B**, **Table S3-4**). Rare subsets were clearly resolved, including innate lymphoid cells 3 (ILC3)^52^ and RORγt+ DCs^53^. These two rare immune populations showed a significant association in our TISME atlas, especially within samples from the colorectal cancer (CRC) and the ovarian cancer (OV) cohorts (**Fig. S8C–D**).

Within the monocyte/macrophage compartment, we identified previously described tumor-associated populations, including the pro-tumoral SPP1-high and the anti-tumoral CXCL9-high tumor-associated macrophages (TAMs)^54,55^ (**Fig. 5D–E**). Although TAMs and monocytes have been extensively characterized using scRNA-seq, much less is known about their chromatin regulation and the key TFs governing their polarization in the TME. We therefore leveraged our multiomic metacell atlas to address this gap by performing multimodal marker analysis that combined TF motif accessibility with TF RNA expression **(Fig. 5F, Table S3 & S5**). CXCL9-high macrophages were strongly marked by both RNA expression and motif accessibility of NFKB and IRF TFs, which are known regulators of the interferon response. Consistent with this, CXCL9-high macrophages displayed the highest score for a previously reported interferon-response signature in monocytes/macrophages^56^ (**Fig. 5G, Table S6**). This signature overlaps with a TAM signature enriched in patients responding to adoptive cell therapy with tumor-infiltrating lymphocytes (ACT-TILs)^57^, which was also significantly up-regulated in CXCL9-high macrophages (**Fig. 5G–H, Table S6**). These findings corroborate our compositional analysis, which showed significant associations between CXCL9-high macrophages and CD8 exhausted T-cell subtypes (**Fig. S8C–D**).

Overall, this extensive metacell-based annotation of a recently published and only partially analyzed multiomic TME atlas highlights how multimodal metacells can accurately resolve highly heterogeneous tissues and deepen our understanding of complex biological systems.

### Multimodal metacell analysis characterizes interferon-primed CD14 monocytes in blood from healthy donors

Given that TAMs can arise from blood monocytes that migrate into tumors and differentiate upon activation, we hypothesized that interferon-primed monocyte populations may already be detectable in PBMCs. To investigate this hypothesis, we first examined the PBMC CITE-seq metacell atlas generated earlier in this study. We identified three monocyte metacell clusters (**Fig. 6A, 4B, S9A**), corresponding to the originally annotated CD14 and CD16 monocytes, as well as an additional small CD14 subtype. This previously unreported CD14 subtype showed clear enrichment for both the CXCL9-high TAM gene signature and a reported interferon-response signature in monocytes/macrophages^56^ (**Fig. 6B, S9B, Table S6**).

**Figure 6:**
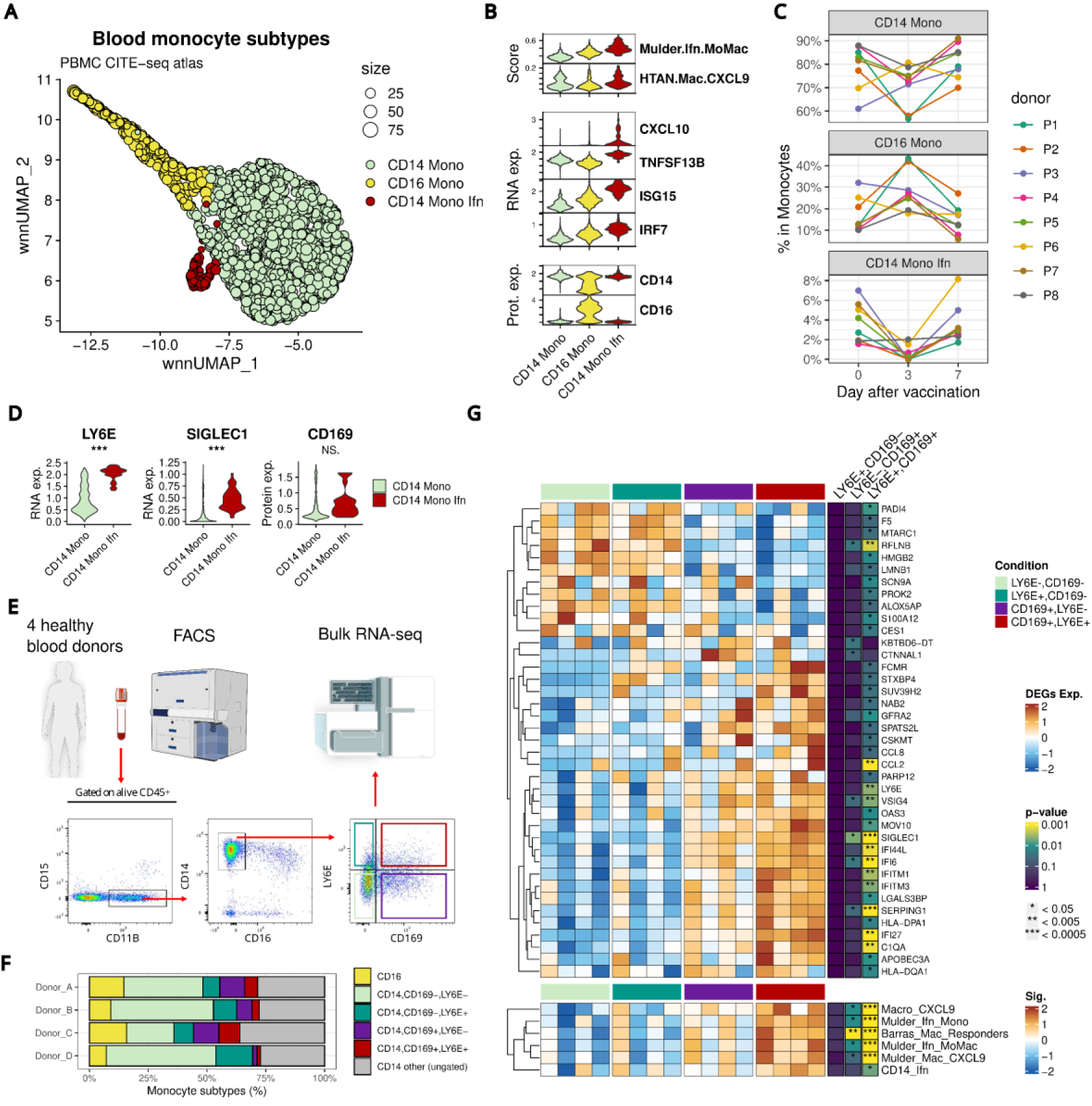
Multimodal metacell analysis characterizes interferon-primed CD14 monocytes in blood from healthy donors. **A** WNN UMAP of monocytes from the CITE-seq PBMC metacells atlas. Metacells are colored by refined blood monocyte subtypes derived from multimodal clustering. **B** Selected Protein and RNA markers as well as signature scores (based on RNA) in the different monocyte subtypes. **C** Proportion of monocyte subtypes at the different time points of the HIV vaccine trial for the 8 donors. **D** Violin plot of *LY6E*, *SIGLEC1* RNA expression and CD169 (encoded by *SIGLEC1*) protein expression in metacells from healthy donors (day 0 of the vaccination trial) in CD14 monocytes compared to interferon-primed CD14 monocyte clusters. (***) indicates an EdgeR FDR < 0.001 and (NS.) indicates a non-significant difference. **E** Experimental workflow implemented to test LY6E and CD169 as surface markers to study interferon-primed CD14 monocytes. A classical FACS strategy was used to gate CD14 monocytes from four healthy blood donors. Four CD14 subpopulations were defined by CD169 and LY6E surface expression, sorted and subjected to bulk RNA-sequencing. **F** Monocyte subtype proportions in each donor were measured by FACS. **G** Upper panel: Heatmap of the protein coding differentially expressed genes between the main CD14+,CD169-,LY6E- monocyte subpopulation and the three other CD14 monocyte subtypes. Lower panel: Selected signature scores differentially expressed between the main CD14+,CD169-,LY6E- monocyte subpopulation and the three other CD14 monocyte subtypes.

The PBMC CITE-seq atlas comprises samples from eight donors collected at three time points in an HIV vaccine trial, enabling us to track monocyte subtype dynamics following vaccination. Interferon-primed CD14 monocytes displayed the most consistent temporal pattern, with a pronounced decrease at Day 3 in all but one donor, followed by recovery by Day 7 (**Fig. 6C**). These findings highlight a potential role for interferon-primed CD14 monocytes in the early immune response and motivated further characterization in healthy individuals.

Differential expression analysis at Day 0 (healthy individuals before vaccination) identified two surface protein-coding genes, *LY6E* and *SIGLEC1*, as significantly upregulated in interferon-primed CD14 monocytes compared with classical CD14 monocytes (**Fig. 6D, S9C, Table S7**). *SIGLEC1* encodes the surface protein CD169, which was included in the CITE-seq antibody panel. Although CD169 protein levels were modestly higher in interferon-primed CD14 monocytes for some donors, no significant differences were observed at the cohort level for CD169 nor any of the proteins in the antibody panel. (**Fig. 6D, S9C**).

We next assessed whether CD169 and LY6E could serve as surface markers to enrich interferon-primed monocytes from fresh blood. We designed a FACS panel and sorted CD14 monocytes into four subpopulations based on CD169 and LY6E expression, using samples from four healthy donors (**Fig. 6E**). All four subpopulations were readily quantified and isolated. As expected, CD14+,CD169–,LY6E– monocytes constituted the major subtype (20–47% of all monocytes per donor; **Fig. 6F, S10A–D and Table S8**). CD14+ monocytes positive for at least one of the two putative interferon-primed markers accounted for 19–26% of all monocytes per donor (**Fig. 6F, S10A–D and Table S8**), slightly higher than the proportion of interferon-primed CD14 monocytes observed in the CITE-seq atlas (**Fig. 6C**).

Bulk RNA-seq of the four sorted populations confirmed upregulation of interferon-related genes, including *SIGLEC1*, *LY6E*, *IFITM1*, *IFITM3*, and *CCL2* in CD14+,CD169+,LY6E+ monocytes compared with the CD14+,CD169–,LY6E– population (**Fig. 6G and Table S9**). Previously reported interferon-response signatures in monocytes/macrophages^56,57^ were most strongly expressed in CD14+,CD169+,LY6E+ monocytes, which also showed the highest expression of the signatures of CXCL9-high TAMs and CD14+ interferon-primed blood monocytes identified in our TISME and PBMC metacell atlases.

Together, these results demonstrate that the combined use of CD169 and LY6E surface markers enables enrichment of interferon-primed CD14 monocytes. This provides a tractable strategy to isolate and study these cells in healthy individuals and suggests that interferon-primed CD14 monocytes, transcriptionally close to anti-tumoral TAMs, may play a key role in early immune responses, as evidenced by their dynamic modulation following vaccination.

## Discussion

In this study, we introduce SuperCell2.0, a framework for identifying robust metacells from single-cell multimodal data, and demonstrate its ability to substantially reduce both dataset size and sparsity in large multimodal single-cell atlases. By aggregating highly similar cells across modalities, our multimodal metacell analysis consistently increased intermodality consistency, as reflected by stronger feature-level correlations between modalities compared with single-cell data. These results support the increasingly common use of coarse-graining strategies in GRN inference from single-cell data24,33,37,58, in which dropout noise and data sparsity remain major limitations^59^.

Building on recent advances in single-cell integration42,44, we incorporated prior knowledge in the form of partial cell-type annotations into the metacell identification process. This semi-supervised strategy reduces metacell impurity and mitigates artifacts that have previously been reported for unsupervised metacell approaches60. Unlike previous metacell approaches relying on a single modality^23,61^ or modality-specific metacells paired post hoc62, SuperCell2.0 further leverages the weighted nearest neighbor (WNN) framework to learn the relative contribution of each modality directly from the data47, thereby identifying more robust multimodal metacells. The SuperCell2.0 framework is also readily extensible to three or more modalities, with minor adaptations of the WNN implementation as previously demonstrated at the single-cell level for Transcriptome-Epitope-ATAC joint sequencing data^63^.

Despite rapid technological progress, single-cell multimodal assays remain substantially more expensive than unimodal profiling. Given the improved intermodality consistency observed at the metacell level, future developments could exploit metacells to integrate modalities profiled independently from the same biological samples. This strategy has already been applied to pair RNA and ATAC data for GRN inference at the metacell level33. Extensions incorporating bridge integration^64^ with a true single-cell multiomic reference, or semi-supervised approaches that leverage partial cell-type annotations could further improve the generation of multimodal metacells from unpaired single-cell datasets. Such advances would be particularly valuable for large public atlases, including HTAN^13,50,51^, where different modalities were often profiled independently for a majority of samples.

Application of the metacell semi-supervised integration workflow to both a PBMC CITE-seq atlas from an HIV vaccine trial and a large 10x Multiome atlas of the tumor microenvironment demonstrated that partial annotations can be effectively leveraged to build large multimodal metacell atlases with fine cell-type resolution. We deliberately used conservative graining levels (10 or 20) to minimize the risk of merging rare or previously unannotated cell populations that may only emerge through large-scale integration. Even at these modest graining levels, the metacell-based workflow yielded substantial reductions in computational cost and atlas size, enabling interactive exploration of datasets that would otherwise be challenging to manipulate at single-cell resolution.

Finally, our extensive metacell-based annotation of the multiomic HTAN atlas illustrates how multimodal metacells can resolve highly heterogeneous tissues and generate new biological insights. Focusing on tumor-associated macrophages (TAMs), we achieved an epigenetic and transcriptional characterization of CXCL9-high TAMs that have recently been identified as predictive biomarkers of immune response and anti-tumor potential^54,57,65,66^. Extending this analysis to a PBMC CITE-seq atlas, we identified an interferon-primed CD14 monocyte subtype that was not reported in the original single-cell study and which displays significant transcriptional similarity to CXCL9-high TAMs, corroborating recent findings from large scRNA-seq atlases of myeloid cells^56^. Importantly, metacell-level marker analysis enabled experimental validation using FACS and bulk RNA-seq, establishing CD169 and LY6E as a combinatorial surface marker strategy to enrich and study these cells in healthy donors. This could help refine further research on this pro-inflammatory monocyte subtype that has been previously analyzed by flow cytometry using CD169 only^67,68^. Our results suggest that interferon-primed CD14 monocytes, transcriptionally related to TAMs with reported anti-tumoral activity, may play a key role in early immune responses. Their dynamic modulation following vaccination that we uncovered in this study supports a model in which these cells rapidly respond to immune challenge, potentially migrating from the bloodstream to the vaccination site or draining lymph nodes, where they may differentiate into interferon-responsive, pro-inflammatory macrophages.

Together, this study introduces a robust approach to facilitate the analysis of large multiomic single-cell datasets based on semi-supervised multimodal metacells. Both in terms of visualisation, analysis and data integration, our results demonstrate the robustness of the metacells built with SuperCell2.0. Our study further highlights how our multimodal metacell frameworks can bridge large-scale data integration with testable biological hypotheses and help characterize rare cell type populations, such as the interferon-primed CD14 monocytes analyzed in this work.

## Methods

### SuperCell2.0 workflow

The unsupervised SuperCell2.0 workflow harnesses up to two modalities of a single-cell experiment to identify robust metacells (**Fig. 1A**). The workflow requires a preprocessed dataset with computed latent spaces for each modality considered. We used dimensionality reduction tailored to each modality: Principal Component Analysis (PCA) was applied to highly variable genes for RNA or proteins for ADT, and Latent Semantic Indexing (LSI) was used for peaks in ATAC data. Next, SuperCell2.0 utilizes these latent spaces to construct a multimodal k-Nearest Neighbor (kNN) graph of the cells using the Weighted Nearest Neighbor (WNN) algorithm implemented via the FindMultimodalNeighbors function in Seurat^47^. By default, in SuperCell2.0, the graph connects each cell to k=30 neighbors, with edge weights reflecting multimodal cell-to-cell affinities computed using the Seurat kernel. In cases where only one modality is provided (e.g., RNA), SuperCell2.0 creates a standard kNN graph based on the corresponding latent space (e.g., PCA), with edge weights corresponding to cell-to-cell affinities computed using a kernel previously proposed for scRNA-seq data^69^. The walktrap algorithm^45^ using weighted random walks is then applied to make a hierarchical clustering of the single cells from the kNN. Metacells are identified from this hierarchical clustering at a specified graining level γ, defined as the ratio of the number of single cells to the desired number of metacells. Single-cell profiles are finally aggregated into metacells by summing raw counts for each modality within each metacell. If single-cell annotations are available, for each of them, SuperCell2.0 annotates a given metacells to the most abundant label within this metacell.

The semi-supervised workflow consists of using a single-cell annotation to build kNN graphs for each cell type separately from the same latent spaces as in the unsupervised workflow (**Fig. 1B**). These graphs are merged with the subset of the global unsupervised kNN graph containing unknown cells and their connections (e.g. edges connecting unknown cells with their neighbors). This new kNN is used as input for metacell identification with the walktrap algorithm. This approach allows us to both preserve connections of unknown cells while avoiding edges between cells annotated to different cell types. In case a complete single-cell annotation is given, it results in a disconnected kNN graph and the identification of pure metacells regarding the given annotation.

SuperCell2.0 is available as an R package and fully integrated into the Seurat v5 framework^64^. Practically, SuperCell2.0 takes as input a preprocessed single-cell Seurat object and outputs a metacell Seurat object at a given γ. After a first run, the hierarchical communities computed with the walktrap algorithm are saved, allowing effortless rescaling of the metacell object at a new γ. The SuperCell2.0 package also provides a comprehensive set of functions for metacell visualization and quality control inside the Seurat v5 framework.

### Preprocessing of single-cell multimodal datasets

The same preprocessing steps as in the original studies were performed with the Seurat v5.1 and Signac^70^ v1.14 R packages for computing single-cell latent spaces used as input for metacell identification (**Table 1**).

#### PBMC 10x Multiome

The PBMC 10x Multiome dataset containing 10,412 annotated nuclei, was retrieved as a Seurat object containing a raw counts matrix for each modality using the SeuratData R package. Cell type labels used for benchmarking and supervision correspond to the Hao et al. annotation obtained from a mapping of the single-nuclei PBMC 10x Multiome RNA data onto the single-cell PBMC CITE-seq atlas followed by manual expert curation^47^. The fragment file was obtained from the 10x website : https://www.10xgenomics.com/datasets/pbmc-from-a-healthy-donor-granulocytes-removed-through-cell-sorting-10-k-1-standard-1-0-0. The joint measurements of RNA and ATAC in single nuclei of human PBMCs were preprocessed as previously47. For ATAC data, MACS2^57^ v2.2.9.1, was used to call narrow peaks in each annotated cell type with the CallPeaks function of Signac (parameters group.by = “seurat_annotations”, and additional.args = “--max-gap 50”). Peaks outside the standard chromosomes or in blacklisted regions of the hg38 genome were discarded. Fragment counts for each called peak were quantified per cell using the FeatureMatrix function in Signac. Cells annotated as “filtered” in the original study were discarded. Dimensionality of the resulting ATAC peak count matrix was then reduced using LSI as previously using a standard Signac workflow. The peak count matrix was normalized using the logarithm of Term Frequency Inverse Document Frequency (TF-IDF) as implemented by Stuart et al.^46^. The TF-IDF matrix was decomposed via SVD to extract the LSI components, and the LSI loadings for each cell were scaled to have a mean of 0 and a standard deviation of 1. For RNA modality, gene expression was normalized and HVGs selected using a regularized negative binomial regression with the SCTransform approach^72^. The top 3000 HVGs were used as input for dimension reduction with a PCA. Latent spaces of the first 40 components of the PCA (RNA) and the LSI (ATAC), excluding the first LSI component which was correlated to ATAC sequencing depth, were retained for metacell identification.

#### BM CITE-seq

The annotated human BM CITE-seq dataset was retrieved as a Seurat object containing a raw counts matrix for each modality using the SeuratData R package. It contains 30,672 high quality and annotated cells with the measurements of 25 ADTs binding to cell surface proteins^46^. Cell type labels used for benchmarking and supervision correspond to the Hao et al. fine cell type annotation obtained from a manual expert annotation of integrated multimodal single-cell data46. Joint measurements of RNA and surface proteins in single-cells of human BM mononuclear cells were preprocessed as previously^46,47^. For RNA modality, gene expression was log normalized and HVGs selected using the standard Seurat workflow. Scaled (centered and reduced) normalized expressions of the top 2000 HVGs were used as input for dimension reduction with a PCA. Protein ADT counts were normalized using a centered log ratio transformation across cells (NormalizeData Seurat function with normalization.method = “CLR” margin = 2), scaled, and used for dimension reduction with a PCA. Latent spaces of the first 30 components of the RNA PCA and the 18 components for the protein ADT PCA were retained for metacell identification.

#### PBMC CITE-seq

We used the published single-cell CITE-seq atlas of human PBMCs consisting of 161,159 single cells with a panel of 208 ADTs47. It gathers measurements for 8 donors profiled at day 0, 3 and 7 of a HIV vaccine trial. RNA and ADT count matrix as well as the multi-level annotation of the atlas was retrieved from the GEO database under the accession number GEO: GSE164378. Joint single-cell measurements of RNA and surface proteins in single cells were preprocessed in each of the 24 samples as previously^47^. For RNA modality, gene expression was normalized and HVGs selected using a regularized negative binomial regression with the SCTransform approach^72^. The top 3000 HVGs were used for dimension reduction with a PCA. Protein ADT counts were normalized in each sample using a centered log ratio transformation across cells (NormalizeData Seurat function with normalization.method = “CLR” and margin = 2) and used for dimension reduction with a PCA. Latent spaces of the first 40 components of the RNA PCA and the first 50 components for the protein ADT PCA were retained for metacell identification in each sample.

#### HSPC 10x Multiome

The annotated human HSPC (CD34+) 10x Multiome dataset^23^, with RNA and ATAC peak counts used in the original study as well as the corresponding fragment file, were retrieved from Zenodo at https://doi.org/10.5281/zenodo.6383269. Joint measurements of RNA and ATAC in single nuclei were preprocessed using a Signac/Seurat workflow similar to the original scanpy workflow. For RNA modality, gene expression was log normalized and scaled (centered and reduced) normalized expression of the top 2000 HVGs were used as input for dimension reduction with a PCA. Dimensionality of the ATAC matrix was reduced using LSI as we did for the PBMC 10x Multiome dataset using a standard Signac workflow (see above). Latent spaces of the first 50 components of the PCA (RNA) and the LSI (ATAC), excluding the first LSI component which was correlated to ATAC sequencing depth, were retained for metacell identification.

#### TISME 10x Multiome

##### In-silico sorting and annotation of tumor-associated immune and stromal cell 10x Multiome data from HTAN

Previously published and annotated 10x Multiome data of colorectal cancer (CRC), pancreatic ductal adenocarcinoma (PDAC), skin cutaneous melanoma (SKCM), uterine corpus endometrial carcinoma (UCEC), ovarian cancer (OV), breast cancer (BRCA), head and neck squamous cell carcinoma (HNSCC) and cervical squamous cell carcinoma (CESC) HTAN cohorts^13,50,51^ were retrieved from the synapse repository (https://www.synapse.org/). Downloaded data consisted in Seurat RNA and ATAC RDS objects, with count matrices as well as fragment files for 122 samples. Each sample was annotated with scATOMIC^41^ v2.0.3 based on raw RNA counts. We used copy number variation (CNV) correction for annotating cancer cells and we assumed one cancer type in each sample (create_summary_matrix scATOMIC function with parameters use_CNV and modify_results set to TRUE). Cells annotated to blood or stromal cell types at the second layer of scATOMIC classification were retained and cancer annotated cells discarded. Cells were subsequently annotated with higher precision as follows. The third layer of scATOMIC classification was used to annotate fibroblast, endothelial, B cells, monocytes/macrophages, plasma, dendritic and mast cells. The fourth layer of scATOMIC classification was used to annotate NK, CD4 and CD8 cells. Cells annotated as unclassified blood cells, unclassified stromal cells or unclassified between two previously annotated cell types were annotated as unknown. This resulting annotation was used to guide metacell identification and batch correction in the semi-supervised workflow of multimodal metacell atlas construction (see below). Cells annotated in the HTAN atlas as tumor, low quality, doublet or tissue specific cells (e.g. non-immune/stromal cell type present in only one cancer type cohort) were discarded. After this filtering, 128,039 cells from 87 samples remained.

##### Generation of a consensus peak set

In each annotated sample, MACS2^71^, was used to call narrow peaks in each annotation of the second layer of scATOMIC (B lymphoid, T/NK lymphoid, myeloid, mast, stromal and unclassified cells) with the CallPeaks function of Signac (parameters group.by = “layer_2”). Peaks outside the standard chromosomes or in blacklist regions of the hg38 genome were discarded. A consensus peak set unifying peaks of all samples was then generated by merging all intersecting peaks (reduce function of the GenomicRanges R package) with a width in between 20 and 10,000 base pairs. To facilitate integration of male and female samples, peaks in the Y chromosome were discarded.

##### Dimension reduction of RNA and ATAC modalities

Joint measurements of RNA and ATAC in single nuclei of immune and stromal tumor-associated cells sorted and annotated with scATOMIC were preprocessed as follows in each of the 87 samples. For RNA modality, raw counts of gene expression were log normalized and HVGs selected using the standard Seurat workflow. The top 2000 HVGs were used as input for dimension reduction with a PCA. For ATAC data, fragment counts for each consensus peak were quantified per cell using the FeatureMatrix function in Signac. Dimensionality of the ATAC matrix was then reduced using LSI as for the PBMC 10x Multiome dataset using a standard Signac workflow (see above). Latent spaces of the first 50 components of the PCA (RNA) and the LSI (ATAC), excluding any LSI component which was correlated (absolute Pearson correlation coefficient above 0.5) to ATAC sequencing depth, were retained for metacell identification.

### Benchmark of metacell identification approaches

#### Metacell approaches and tested scenarios

The different methods were tested on the preprocessed (low quality cell filtered) PBMC 10x Multiome and BM CITE-seq datasets at γ=75 and γ=20. The unsupervised multimodal workflow of SuperCell2.0 was compared to SEACells v0.3.3 (Persad et al.) using RNA or ATAC/Protein and MetaCell2 (v0.9.0) using RNA. The unsupervised unimodal workflow of SuperCell2.0 was also tested using RNA or ATAC/Protein. SEACells^23^ and MetaCell2^57^ were implemented using their default parameters and author guidelines. The same latent spaces (e.g. same components) for each dataset and each modality were used as input for SuperCell2.0 and SEACells while MetaCell2 was given raw counts RNA matrix of each dataset. Following the author’s recommendation, some genes, including mitochondrial and ribosomal protein genes, were not used for metacell identification with MetaCell2.

The semi-supervised multimodal workflow of SuperCell2.0 was tested as the unsupervised one on the PBMC 10x Multiome and BM CITE-seq dataset at γ=75 using as input 0, 25, 50, 75 and 100 % of annotated cells (according to original dataset annotation).

We referred as scenarios the pairs of dataset and γ tested.

#### Coverage plots

Single-cells WNN UMAP of the PBMC 10x Multiome, BM CITE-seq, were computed using the same latent spaces as for the metacell identification. First, the FindMultimodalNearestNeighbor Seurat function (with the default k=20 value) was used to build a WNN graph using RNA and Protein or ATAC latent spaces which was then used as input for the RunUMAP function of Seurat (parameter nn.name = “weighted.nn”). Single-cell WNN UMAP coordinates were then averaged in each metacells to make the metacell coverage plots^24^ for each tested scenario. Coverage plots were also performed for the sample P1 on day 0 of the PBMC CITE-seq atlas and the HSPC 10x Multiome data with the multimodal SuperCell2.0 unsupervised workflow at γ=75 for the intermodality consistency analysis.

#### Purity

The purity of a metacell *m* according to a single-cell annotation is defined as the proportion of cells from the most abundant cell type of the given annotation in this metacell^24^:

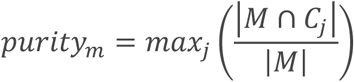

With *M* the set of cells belonging to the metacell *m* and *C_j_* the set of cells belonging to cell type *j*.

Metacell purities according to the original dataset annotation were computed for each tested scenario.

#### Compactness and separation

We used the previously introduced metrics of metacell compactness and separation to evaluate metacell approaches^23^. Following their original definition, compactness and separation were computed for each modality in the diffusion map derived from the latent space used for metacell identification (e.g. PCA for RNA and Protein, LSI for ATAC). For each latent space in each dataset, *N* = 10 diffusion components were computed as previously using the palantir v1.3.3 python package^69^.

The compactness of a metacell *m* corresponds to its homogeneity and is defined as the average variance of single cell coordinates within a metacell (the lower the better).

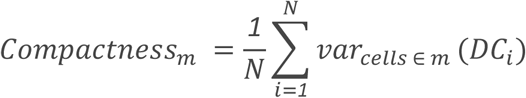

where *DC* is the matrix of diffusion components computed from single-cell data for a given modality.

The separation of a metacell *m* represents its distance from the closest other metacell *l*. It is defined as the Euclidean distance between centroids of the single-cell coordinates forming the metacells in the diffusion space of a given modality (the higher the better).

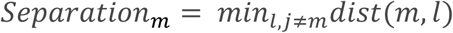

Compactnesses (separations) of a metacell *m* in a given modality were scaled from 0, the lowest, to 1, the highest across all values obtained with the different metacell approaches tested in a given scenario:

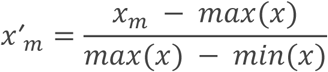

where *x* is the compactness or separation vector of values from all tested metacell approaches for a given modality in a given scenario and *x′* the corresponding vector of scaled values.

This allowed us to define the multimodal compactness (separation) *y_m_* of a metacell *m* as the average of the scaled compactnesses (separations) of the metacell *m* for the *M* different modalities of the dataset for a given scenario:

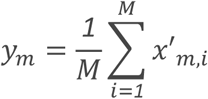

where *x^′^_m,i_* is the scaled compactness (or separation) of metacell *m* in the modality *i*.

#### Computational load

Metacell identification and prior single-cell preprocessing for each tested scenario were included in a Snakemake pipeline^73^. The computational load of each metacell tool in each unsupervised scenario was assessed according to CPU time and peak memory use. The Snakemake benchmarking function was used to measure time and peak memory usage (max PSS). For MetaCell2 scenarios that take a single-cell RNA count matrix as input, CPU time and peak memory usage were assessed directly on metacell identification. For SEACell and SuperCell2.0 scenarios, computational load includes both metacell identification and the required preprocessing of the modality or modalities used as input. In these scenarios, CPU time corresponds to the sum of CPU times of modality preprocessing and metacell identification, and peak memory usage corresponds to the highest peak memory usage measured between preprocessing and metacell identification. The benchmarking was performed on one node of an HPC cluster with 512G of RAM.

### Analysis of intermodality consistency at the metacell level

We analyzed intermodality consistency in metacells obtained with the unsupervised and multimodal workflow of SuperCell2.0 through intermodality feature correlations at increasing γ from 10 to 200. For CITE-seq data, we conducted this analysis on the BM and the sample P1 at time 0 (“P1_0”) of the HSPC atlas. For 10x Multiome data we implemented this analysis on the PBMC 10x and HSPC Multiome datasets.

#### Analysis of gene and transcription factor motif accessibilities

For the two 10x Multiome datasets, at each tested γ, fragments were aggregated at the metacell level in order to compute gene and TF motif chromatin accessibilities at the metacell level. Gene activity representing gene chromatin accessibility was computed using the GeneActivity function of the Signac package which sums fragment counts falling into the gene body and the promoter region (from -2000 bp of the TSS to the Transcriptional terminal site). Gene activities were computed for the top 4000 HVGs at the RNA level for which a gene activity could be computed (gene width of at most 50 kbp). TF motif accessibilities were computed using chromVAR^74^ v1.24.0 using the Hocomoco TF motif database75.

#### Analysis of intermodality correlations

Correlations between RNA gene expression and corresponding surface protein abundance in CITE-seq data as well as correlation between gene activity (ATAC) and RNA gene expression for 10x Multiome data were assessed at the different γ taking into account metacell sizes by computing weighted Pearson correlation using the weights R package^76^ v1.0.4. As controls, correlation tests were also performed at each γ, in random metacells (randomly grouped single-cells) and in subsampled single-cells. For subsampling at high γ, we set the r correlation to zero when insufficient cells expressed both features to calculate it.

In single-cell 10x Multiome data, we identified the TFs marking a main cell type (B cells, T cells, Dendritic cells, Monocytes) both by a significant (p-adjusted value of wilcoxon test below 0.05) increase of their motif accessibility (mean difference value between chromVAR z-score” above 0.1) and increase of their gene expression at the RNA level (log fold change of gene expression above 0.1) using the wilcoxauc function from the presto package v1.0.0. For this subset of TFs, weighted correlations between their motif accessibility and their RNA gene expression were assessed as for the previous intermodality feature correlations.

#### Gene regulatory network inference

We used Pando v1.1.1 to infer regulons33, modules of a TF and its target genes in the PBMC 10x Multiome dataset at each γ. Pando models the expression of target genes as functions of TF RNA expression - binding site accessibility pairs using generalized linear models (GLM).

At each γ, the metacell ATAC data were normalized using the TF-IDF method by Cusanovitch et al.^78^ (using the RunTFIDF function from Signac with method=2) and metacell RNA were log normalized. Pando was run on the normalized metacell data using as input target genes the top 4000 HVGs identified at the single-cell RNA level (genes selected with the FindVariableFeatures function of Seurat). Candidate regulatory elements were identified by intersecting ATAC peaks with TF motifs and with target genes. We modified the Pando procedure to consider metacell sizes. First, we included the size of the metacells in the evaluation of correlations between peaks accessibility and gene expression (using wtd.cor from the weights R package v1.0.4). Second, we included the metacell size in the GLMs as weights in the svyglm function from the survey R package (v4.4-2) to model the expression of the target genes. GRN inference was performed using the infer_grn function from the Pando package (with parameter peak_to_gene_method = “GREAT” and the other default parameters). Highest significant weights of the GLMs for each target gene were then collected to build regulons (using the find_modules.GRNData function from Pando with the following parameters: p_thresh=0.05, nvar_thresh=2, min_genes_per_module = 1, rsq_thresh = 0.05). For each of them, enrichment in known targets from the CollecTRI database^49^ was assessed using a hypergeometric test.

### Metacell multimodal atlas construction

The PBMC CITE-seq and the TISME 10x Multiome altases were constructed as follows (**Fig 4A**). After datasets pre-processing to obtain a single-cell latent space for each modality for each sample (see above), metacells were identified per sample using the SuperCell2.0 semi-supervised and multimodal workflow. Then, semi-supervised batch correction at the metacell level was performed in each modality using anchor-based reciprocal PCA integration with STACAS^42^ (reciprocal LSI with Signac for ATAC). This method identifies biologically similar cells across datasets termed anchors, by detecting mutual nearest neighbors, which serve as correspondences between datasets and form the basis for estimating and correcting batch effects. STACAS uses the prior knowledge of cell labels to discard anchors between cells annotated to different cell types to supervise batch correction. Batch corrected metacell latent spaces of each modality were eventually integrated in a WNN graph that we used as input for UMAP visualization and multimodal metacell clustering. Metacell atlases were also constructed with a similar but fully unsupervised workflow (without any usage of cell annotation for metacell identification and batch correction) to evaluate the benefit of prior knowledge usage.

#### PBMC CITE-seq metacell atlas

We used γ=20 for the PBMC CITE-seq samples and the previously published annotations to guide SuperCell2.0 and STACAS. More precisely we used the first annotation level of Hao et al.^47^ comprising B, CD4 T, CD8 T, NK, Monocytes, DC, other T and other cells. For the two latter we used the second level of annotation dividing “other T” into double negative (dn)T, gamma-delta (gd)T and mucosal-associated invariant T cells (MAI)T cells and “other” into HSPC, ILC and erythrocytes. We took the hyperparameters used in the original study for our metacell integration^47^. We selected the top 2000 HVG across samples at the metacell level as RNA integration features using the SelectIntegrationFeatures function of Seurat and all features (208) were used for ADT integration. We used the unscaled log Normalize expression for RNA batch correction. Metacell ADT counts were normalized using CLR normalization and scaled for ADT batch correction. We used the eight samples at day 0 before vaccination as reference samples for STACAS in order to limit the number of anchor searches between pairs of dataset and reduce the associated computational burden. As in the original study at single-cell level, we took the first 40 components for RNA batch correction and the first 50 components for ADT batch correction.

The resulting RNA and ADT corrected metacell latent spaces by STACAS were integrated in a WNN graph of metacells using the Seurat function FindMultimodalNeighbors. We then made a UMAP visualization of the metacell multimodal atlas from this WNN graph (WNN UMAP) using the RunUMAP Seurat function (min.dist = 0.2). Finally, a Weighted Shared Nearest Neighbors (WSNN) graph was derived from the WNN graph of metacells and clustered with the Smart Local Moving (SLM) algorithm scanning 10 resolutions from 0.5 to 1.5 using the FindClusters Seurat function.

We used the same workflow to build the PBMC CITE-seq atlas with decreasing amount of annotated cells in order to evaluate the ability of a semi-supervised workflow to leverage priori knowledge. To do so we randomly hid 5%, 10% and 50% of input cell labels.

#### TISME 10x Multiome atlas

We used a target γ of 10 in order to have at least 200 metacells in each sample of the TISME 10x Multiome atlas, i.e. γ = min([10,n_cells_smp/200]) and used the scATOMIC annotations (see above) as input labels for SuperCell2.0 and STACAS. We defined 16 reference samples for the integration by selecting for the 8 cancer cohorts the two datasets, presenting the highest number of different cell types based on scATOMIC annotations. For RNA, as previously for the PBMC CITE-seq atlas, we selected the top 2000 HVG across samples at the metacell level as integration features and used their log-normalized and unscaled expressions for batch correction. To do so, we used reciprocal PCA anchor-based integration using the first 50 components with STACAS.

For ATAC, we selected the top 75% of most accessible peaks across the whole metacell atlas as integration features and all peaks were normalized using the Cusanovich & Hill TF-IDF normalization^78^ (*TF* × *log*(*IDF*)), as we found that it was better at separating scATOMIC labels at the metacell levels than the Stuart & Butler TF-IDF implementation^46^ (*log*(*TF* × *IDF*)). To correct batch effects in the ATAC modality of our metacells, we implemented a similar semi-supervised workflow as STACAS for RNA, using anchor-based reciprocal LSI integration in the Seurat/Signac R framework. Briefly, we first computed an uncorrected LSI latent space of our whole metacell atlas as well as LSI latent spaces for each metacell sample keeping the first 50 components and discarding all components with an absolute Pearson correlation coefficient with ATAC sequencing depth above 0.5. Integration anchors between samples were found using reciprocal LSI projection, thanks to the FindIntegrationAnchors Seurat function (reduction = “rlsi”). We used the same 16 reference samples as for the RNA batch correction. We then filtered anchors between cells annotated as different cell types by scATOMIC before using them to correct the initial LSI atlas latent space thanks to the IntegrateEmbeddings function of Seurat. To ensure stability, the k.weight parameter was iteratively optimized upon integration failure, decrementing until convergence.

RNA and ATAC batch corrected latent spaces of metacells were integrated in a WNN graph using the FindMultimodalNeighbors Seurat function. We then made a UMAP visualization of the metacell multimodal atlas from this WNN graph (WNN UMAP) using the RunUMAP Seurat function (min.dist = 0.2). Finally, a Weighted Shared Nearest Neighbors (WSNN) graph was derived from the WNN graph of metacells and clustered with the Smart Local Moving (SLM) algorithm^79^ scanning 10 resolutions from 0.5 to 1.5 using the FindClusters Seurat function.

### Benchmark of metacell multimodal atlas integration

#### Metacell purity

We evaluated metacell identification of the (semi-)supervised compared to unsupervised metacell workflows by comparing metacell purity distribution for the different annotations. For the PBMC CITE-seq atlas we computed metacell purities for the main cell type annotation, used as input prior knowledge (see above), and for the fine cell type annotation, named “celltype.l2” in the published metadata^47^. For the TISME 10x Multiome atlas we used the scATOMIC annotations, used as input prior knowledge (see above), and HTAN published annotation^13^.

#### Batch correction

We evaluated batch correction efficiency of the (semi-)supervised compared to unsupervised metacell workflows using cell type Average Silhouette Width (ASW), measuring cell type separation and Cell type integrative Local Inverse Simpson Index (CiLISI), measuring batch mixing in a cell type aware manner. We chose these two metrics as it has been shown that they correctly recapitulate bio-conservation and batch mixing scores made of an exhaustive panel of single-cell integration metrics^42^. Metacells were split into pure metacells according to the cell type labels and we developed weighted implementations of these metrics to take into account the purity and size of the metacells.

For a metacell *i* composed of cells annotated to *k* different cell types, we considered the corresponding *k* pure sub-metacells of sizes *w*. Sub-metacells of unknown cell type were discarded.

We computed the weighted silhouettes *s* of a sub-metacells *j* as we previously described^21^. We obtained the overall cell type ASW for a given atlas of N metacells using a sample-weighted mean of the sub-metacells *s* :

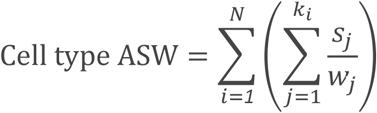

We computed the CiLISI *l* of a sub-metacells *j* as previously described for single-cells^42^. We obtained the overall CiLISI for a given atlas of *N* metacells using a sample-weighted mean of the sub-metacells *l* :

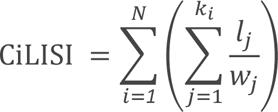

Cell type ASW and CiLISI were computed using Euclidean distance in the metacell corrected latent space of each modality obtained with the (semi)-supervised and the unsupervised workflows as well as for the uncorrected latent space. Cell type ASW and CiLISI were computed using the 2 different annotations for each atlas as for metacell purity (see above).

#### Computational load

The computational load of the (semi-)supervised and the unsupervised metacell workflows, from preprocessing of single-cell samples to final WNN graph construction and clustering was assessed for the two metacell atlases according to CPU time and peak memory usage (max PSS) measured by the Snakemake benchmarking function. The computational load of building equivalent single-cell atlases was also benchmarked. This single-cell workflow consisted in the original workflow of Hao et al. for the PBMC CITE-seq atlas, i.e. Seurat anchor-based unsupervised workflow^47^. For the TISME 10x Multiome atlas, we benchmarked a single-cell semi-supervised workflow similar to the one we developed for metacells (without prior metacell identification per sample).

The benchmarking was performed on one node of an HPC cluster with 512G of RAM. 6 cores were used to parallelize sample preprocessing both in the single-cell and the metacell workflows for the benchmark of the TISME 10x Multiome atlas while no parallelization was used for the PBMC CITE seq atlas.

### Metacell atlas annotations

All available single-cell annotations were aggregated at the metacell level with SuperCell2.0 which assigned a given metacells to the most abundant label within this metacell.

#### Monocyte subtypes in the PBMC CITE-seq atlas

Using the results of metacell multimodal clustering at a resolution of 0.8 we identified an additional cluster of CD14 monocytes with an interferon-response signature (CD14_Mono_Ifn in addition to CD14_Mono and CD16_Mono).

#### Annotation of the TISME 10x Multiome atlas

Starting from the results of the multimodal clustering at a resolution of 1.2 we first identified 26 metacell clusters. We investigated them one by one, analyzing their composition in single-cell scATOMIC and HTAN annotations and looking at their expression for known markers of cell type from the TME both at the gene expression (RNA) and gene activity (ATAC) levels. We could annotate 19 of them to fine cell types of the immune and stromal TME. For the remaining 7 metacell clusters we found that subclustering using the FindSubCluster function of Seurat improved cluster homogeneity regarding single cell annotation and relevance of gene expression/activity markers. More precisely, subclustering of cluster 3 identified the different CD8 T subtypes, cluster 4 could be divided into Monocytes subtypes, cluster 6 into CD4 T subtypes, cluster 7 into Fibroblast subtypes, cluster 8 into Endothelial subtypes, cluster 14 into cycling CD4 T and cycling CD8 T cells and cluster 16 into DC subtypes. In total, our clustering analysis revealed 40 fine clusters that we placed into 8 major types: Endothelial, Fibroblast, Mural, DC, Monocytes/Macrophages, Plasma, B, T, NK and Mast cells.

Despite the initial removal of single-cells annotated as doublets in the published HTAN annotation, 5 fine clusters presented mixed HTAN and/or scATOMIC labels suggesting potential doublets: Fibroblast/Endothelial doublets for cluster 8_1, Fibroblast/T cells doublets for cluster 7_0_1, Fibroblast/Plasma cells doublets for cluster 7_2, T cells/Macrophages doublets for cluster 3_3 and Plasma cells/Macrophages doublets for cluster 22. We confirmed the nature of these doublets by computing major type signatures at the metacell level with the AddModuleScore function of Seurat, using as input markers of major types that we identified using a pseudobulk differential gene expression analysis (see below). To confirm the single-cell origin and exclude a metacell impurity origin of these doublets, we also computed these signatures (using the same markers) on single cells. Metacell clusters of doublets were discarded from the final WNN UMAP representation of the atlas and downstream analysis.

### Marker and signature analysis in metacell atlases

#### Pseudobulk differential expression analyses

To identify cluster markers across RNA, gene activity, and surface protein expression we performed pseudobulk differential expression analyses using edgeR v4^80^. For each assay, raw counts were aggregated by sample and cell type cluster, and low-quality pseudobulk samples (library size <50,000 for TISME atlas and library size <40,000 for the CITE=-seq atlas) were excluded. Feature-level filtering was performed based on minimum expression thresholds across clusters (e.g., CPM ≥10 in ≥70% of samples within the smallest cluster group for RNA and gene activity and CPM ≥1 in ≥70% of samples within the smallest cluster group for ATAC). After Trimmed Mean of M-values normalization, a design matrix encoding cell type clusters and the estimateDisp function from edgeR were used to estimate dispersions robustly (robust = TRUE). A negative binomial generalized log-linear model was fitted to the read counts for each gene using the glmQLFit function from edgeR (robust = TRUE).

We applied quasi-likelihood F-tests to test each cluster against all others using a contrast matrix approach. Differentially expressed features were defined using an FDR threshold of 0.05. Only upregulated markers (logFC >0) were retained.

#### Multimodal markers of cell types in the TISME 10x Multiome atlas

First, markers were identified for each major type (excluding clusters of doublets) both at the gene expression (RNA) and gene activity (ATAC) level, testing each major type against all the others. Second, in each major type (e.g. Monocytes/Macrophages), subtype (e.g. CXCL9-high TAMs) markers were obtained by testing each subtype against all the other subtypes composing the major type. T and NK subtypes were analyzed together at this level. We drew volcano plots (−*log*10*FDR* = *f*(*logFC*)) from the unfiltered results of the differentially expressed gene analysis of CXCL9-high TAMs (**Fig. S9B**).

#### Multimodal TF markers of tumor-associated monocytes/macrophages

ChromVar TF motif accessibility scores of tumor-associated monocyte and macrophage metacells were computed as previously for PBMC and HSPC 10x Multiome data. These scores were then averaged in each subtype for each sample and used to identify TF motif markers of subtypes using Wilcoxon tests (one subtype versus all others). Multimodal TF markers of tumor-associated monocytes/macrophages were finally retained as transcription factors for which both RNA expression and Motif accessibility is significantly upregulated (FDR below 0.05) in one subtypes compared to other tumor-associated monocytes/macrophages.

#### Peripheral Monocytes in the PBMC CITE-seq atlas

Markers of peripheral monocyte subtypes were identified using the same pseudobulk approach both at the gene expression (RNA) and surface protein expression (ADT) levels. The donor variable was considered in the model as a covariable. This analysis was performed only at day 0 before vaccination and using time from vaccination as a covariable in EdgeR’s model.

#### Transcriptomic signatures scoring

Tumor-associated macrophage/monocyte metacells from the TISME 10x Multiome atlas and peripheral monocytes metacells from the PBMC CITE-seq atlas were scored for signatures of interferon response established from a published study of a single-cell atlas^56^. From the DEG results of this study, we defined the Mulder_Ifn_Mono signatures, corresponding to genes significantly upregulated (p-adjusted value < 0.05 & LogFC > 0.25) by a cluster of interferon-responding monocytes, Mulder_Mac_CXCL9 to genes upregulated by a cluster of interferon-responding macrophages and Mulder_Ifn_MoMac to the intersection of the two latter (**table S6**). Tumor-associated macrophage/monocyte metacells were also scored for Barras_Mac_Responders, a previously published signature of tumor-associated macrophage response to ACT-TILs^57^. Peripheral monocyte metacells were also scored for TISME_CXCL9_TAM, the signatures of the CXCL9-high TAM subtypes by using their marker genes that we had identified at the RNA level in the TISME atlas. All signatures were computed using the AddModuleScore function of Seurat.

### Compositional analysis in metacell atlases

#### Immune cell type associations in the TME

We analyzed correlations of T/NK with myeloid (Monocytes/Macrophages, DCs) subtypes abundances in the TISME 10x Multiome atlas. To do so we retained only samples with at least 100 single cells in metacells that we annotated to T/NK or myeloid subtypes. Then, for the 79 remaining samples, we computed the proportion of each T/NK (myeloid) subtypes in the T/NK (myeloid) population at the single-cell level. Finally, we computed the Pearson correlation coefficient for each pair of T/NK - myeloid subtypes. We visualized significant correlations (p-value of Pearson correlation test below 0.05) in a matrix using the corrplot R package^81^ (**Fig. S8C**).

#### Peripheral monocytes subtypes dynamics during HIV vaccine trial

We analysed variation in monocyte composition during the vaccine trial in the PBMC CITE-seq atlas by computing the percentages of monocyte subtypes with respect to the whole monocyte population at the three different time points of the vaccine trial in each of the 8 donors (**Fig. 6C**).

### Analysis of monocytes from fresh blood of healthy donors

#### Sorting of monocytes and RNA extraction

The processing of blood from healthy donors was approved by the Commission cantonale d’éthique de la recherche sur l’être humain (CER-VD, protocol 2024-02022). Fresh peripheral blood from healthy donors was obtained from the Centre Transfusion Interrégionale CRS (Epalinges, Switzerland) and processed within 2 hours of collection. Whole blood was directly incubated in 1× RBC lysis buffer, incubated at room temperature for 10 minutes, and centrifuged at 400 × g for 5 minutes. The next steps of the staining and sorting were performed at 4°C. This procedure was repeated for a total of two rounds of RBC lysis. The resulting cell pellets were resuspended in FACS buffer (PBS + 0.5% BSA (Jackson ImmunoResearch, 001-000-162) + 2 mM EDTA (Invitrogen, 15575-038)) and transferred to 96-well plates for staining, followed by one wash step. Cells were blocked with human TrueStain FcX (BioLegend, 422302) for 20 minutes and subsequently stained in 50 µL/million cells of Brilliant Stain Buffer (25% v/v in FACS buffer, BD Horizon, 566349) with antibodies against CD169 (PE, BioLegend, 346003, 1:100), CD45 (PE/Cy7, BioLegend, 304015, 1:200), CD14 (FITC, BioLegend, 325604, 1:200), CD11b (BV421, BioLegend, 101236, 1:200), CD15 (BV510, BioLegend, 323028, 1:200), CD16 (BV785, BioLegend, 302045, 1:200), and LY6E (APC, Creative Biolabs, MOB-636, 1:100). Fluorescence minus one (FMO) controls for CD169 and LY6E were used to define gates, allowing ∼0.1% false-positive events in the respective FMO controls. Immediately prior to sorting, DAPI was added for live/dead discrimination. Cells were sorted on a FACSAria II cell sorter (BD Biosciences) directly into 350 µL RLT buffer supplemented with DTT (Sigma-Aldrich, 1003440138) and stored at −80 °C until RNA extraction using the RNeasy Micro Kit (Qiagen, 74104) with on-column DNase digestion.

#### Library preparation and sequencing

RNA quality was assessed on a Fragment Analyzer (Agilent Technologies). The RNAs had RQNs between 7.4 and 9.2. RNA-seq libraries were prepared from 1 ng of total RNA with the Takara SMART-seq mRNA LP kit (Takara Bio Inc., 634768). Libraries were quantified by a fluorometric method (QubIT, Life Technologies) and their quality assessed on a Fragment Analyzer (Agilent Technologies).

Sequencing was performed on an Element Biosciences Aviti sequencer with a 2×75 cycles Cloudbreak FS High Output sequencing kit (run as single read 150). Base calling and demultiplexing were done with bases2fastq version: 2.0 with filter mask R1:N15Y15N*.

#### Analysis of the monocyte RNA-seq data

The raw data (fastq files) were preprocessed using the nf-core RNA-Seq pipeline^82^ (version 3.14.0). The human hg38 reference genome and its associated annotations were retrieved from ensembl (release 115). The nf-core pipeline was run with the following parameters: --aligner star_salmon –--extra_salmon_quant_args ‘--fldMean 320 --fldSD 30 --seqBias --gcBias’. Hence, STAR^83^ (v2.7.9a) was used to map the reads to the hg38 reference genome and assign the alignments to transcripts. Salmon^84^ (v1.10.1) was then used to perform BAM-level quantification. Gene-level counts were retrieved using the tximport^85^ R package (v1.34.0) and lowly expressed genes were filtered out using the filterByExpr function from the edgeR R package (v4.4.0)^80^. Normalization factors were computed using edgeR’s trimmed mean of M-values (TMM) method (normLibSizes function) and gene dispersions were estimated with the estimateDisp function. A quasi-likelihood negative binomial generalized linear model was fitted for each gene with the glmQLFit function from edgeR. We applied quasi-likelihood F-tests to perform pairwise comparisons and identify genes significantly differentially expressed in at least one of the comparisons. The Gene Set Variation Analysis (GSVA)^86^ R package (v2.0.0) was used to compute GSVA enrichment scores for each sample and different gene sets previously considered in the paper, i.e. signatures of interferon response established from a published study of a single-cell atlas (Mulder_Ifn_Mono, Mulder_Ifn_MoMac, Mulder_Mac_CXCL9)^56^, a previously published signature of tumor associated macrophages response to ACT-TILs (Barras_Mac_Responders)^57^, a signature (Macro_CXCL9) characteristic of the CXCL9-high TAM subtype identified in the TISME 10x Multiome atlas and a signature (CD14_Ifn) characteristic of the CD14_Ifn subtype identified in the PBMC CITE-Seq atlas.

### Statistics

Statistics were computed with R software v4.3.3. The different statistical tests used are detailed in the specific methods sections above and in the figure legends.

## Data availability

Cell annotations of the PBMC CITE-seq as well as the TISME 10x Multiome metacell atlases are available on Zenodo at https://doi.org/10.5281/zenodo.18613560. This Zenodo repository also contains the gene count matrix of the monocyte RNA-seq data, as well as the containers packaging the software environments of this study (SIF files).

## Code availability

SuperCell2.0 is available as an R package on github at https://github.com/GfellerLab/SuperCell/tree/supercell-2.0. The Snakemake pipeline used to build and analyze the immune and stromal TME metacell atlas is available at https://github.com/GfellerLab/SuperCellMultiomicsHTAN. The Snakemake pipeline used for all the other analyses is available at https://github.com/GfellerLab/SuperCellMultiomicsAnalyses.

## Acknowledgements

We acknowledge the Swiss National Science Foundation (SNSF) Project Grant (310030_197514) (DG), SNSF Advanced Grant TMAG-3_209224 (JAJ), and the Ludwig Institute for Cancer Research (JAJ) for funding support. Benoît Duc was supported by an MD-PhD fellowship from the Fondation ISREC. Bastien Dolfi was supported by a Momentum Fellowship from The Mark Foundation for Cancer Research. We warmly thank Danny Labes and Mariela Castelblanco from the UNIL-Agora flow cytometry platform as well as Johann Weber, Julien Marquis and Corinne Peter from the Lausanne Genomic Technologies Facility (GTF) for their technical support. We thank Mariia Bilous, Santiago J. Carmona and Massimo Andreatta for valuable conversations related to this manuscript.

## Notes

https://github.com/GfellerLab/SuperCell/tree/supercell-2.0

https://github.com/GfellerLab/SuperCellMultiomicsAnalyses

https://github.com/GfellerLab/SuperCellMultiomicsHTAN

https://zenodo.org/records/18613560

### Competing Interest Statement

The authors have declared no competing interest.

