## Supplementary Figures for "SuperCell2.0 enables semi-supervised construction of multimodal metacell atlases"

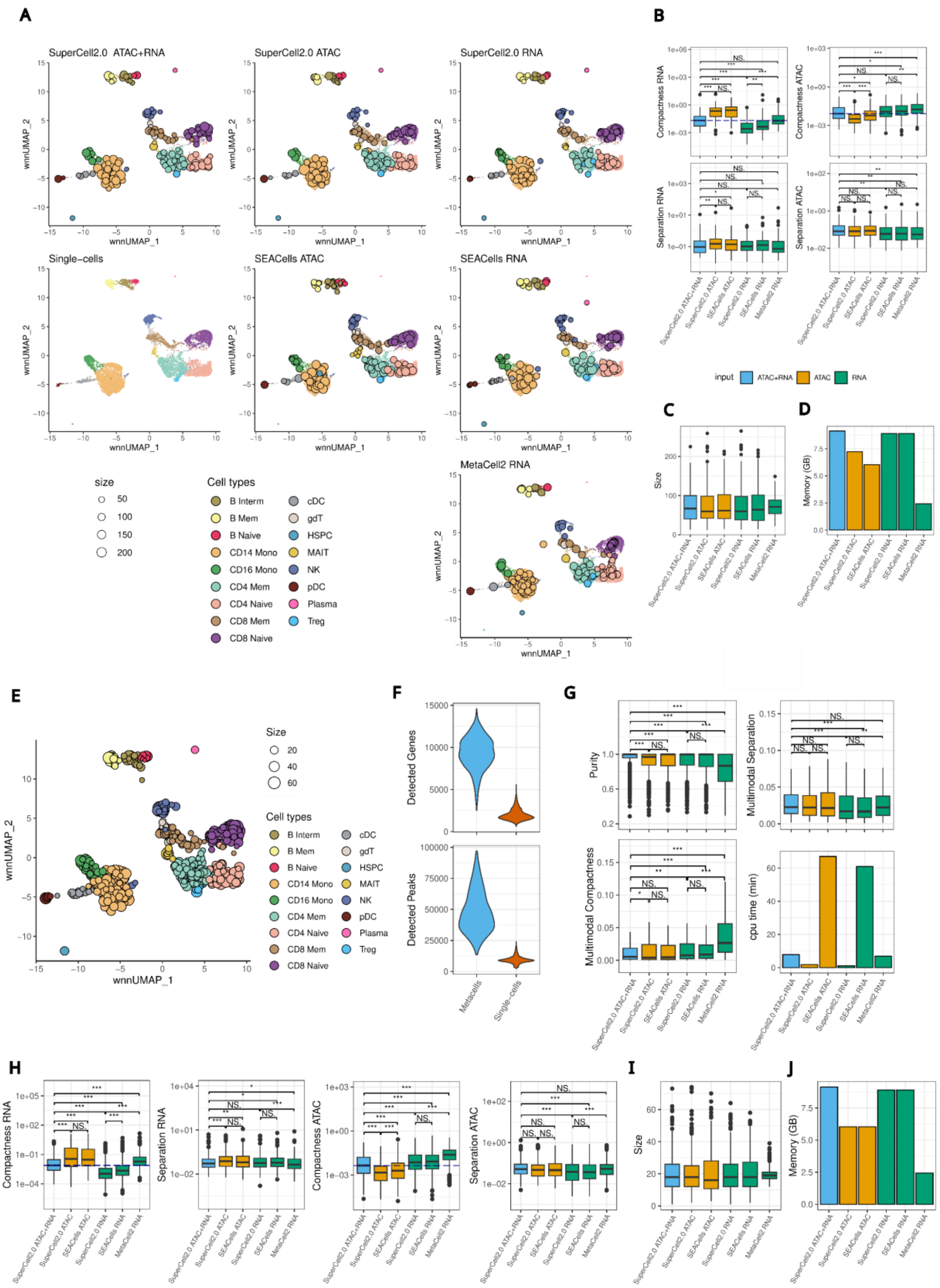


**Supplementary Figure 1: Benchmark of the unsupervised workflow of SuperCell2.0 on PBMC 10x Multiome data. A** Coverage plots of the single-cell WNN UMAP of the metacells identified with SuperCell2.0, SEACells and MetaCell2 using as input ATAC with RNA, ATAC only and RNA only (γ=75). **B** RNA and ATAC compactnesses and separations (log scale) for metacells obtained with SuperCell2.0 (unsupervised), SEACells and MetaCell2 using as input ATAC &

RNA, ATAC only and RNA only. The horizontal dashed lines in compactness boxplots correspond to the median of the metacell compactnesses obtained with SuperCell2.0 multimodal workflow. **C** Size distributions of the corresponding metacells. To note that MetaCell2 algorithm keeps metacell size between a range. **D** Memory peak usage of metacell identification for SuperCell2.0, SEACells and MetaCell2 using as input ATAC & RNA, ATAC only and RNA only. **E** Coverage plot of the single-cell WNN UMAP of metacells identified at $\gamma$=20 with the multimodal workflow of SuperCell2.0. **F** Detected gene and peak count distributions of the metacells in **E** and single cells for the PBMC 10x Multiome data. **G** Metacell purities, multimodal separations and compactnesses (outliers not shown), and CPU time of the tools SuperCell2.0, SEACells and MetaCell2 using as input ATAC & RNA, ATAC only and RNA only ($\gamma$=20). **H** RNA and ATAC compactnesses and separations of the tools SuperCell2.0, SEACells and MetaCell2 using as input ATAC & RNA, ATAC only and RNA only ($\gamma$=20). The horizontal dashed lines in compactness boxplots correspond to the median of the metacell compactnesses obtained with SuperCell2.0 multimodal workflow. **I** Metacell sizes and **J** Memory peak usage of the tools SuperCell2.0, SEACells and MetaCell2 using as input ATAC & RNA, ATAC only and RNA only ($\gamma$=20). Differences in metric distributions were tested using Wilcoxon tests in **B**, **G** and **H** with results grouped into (***) for p-values < 0.001, (**) for p-values < 0.01, (*) for p-values < 0.05 and no significant difference (NS.) for p-value > 0.05.

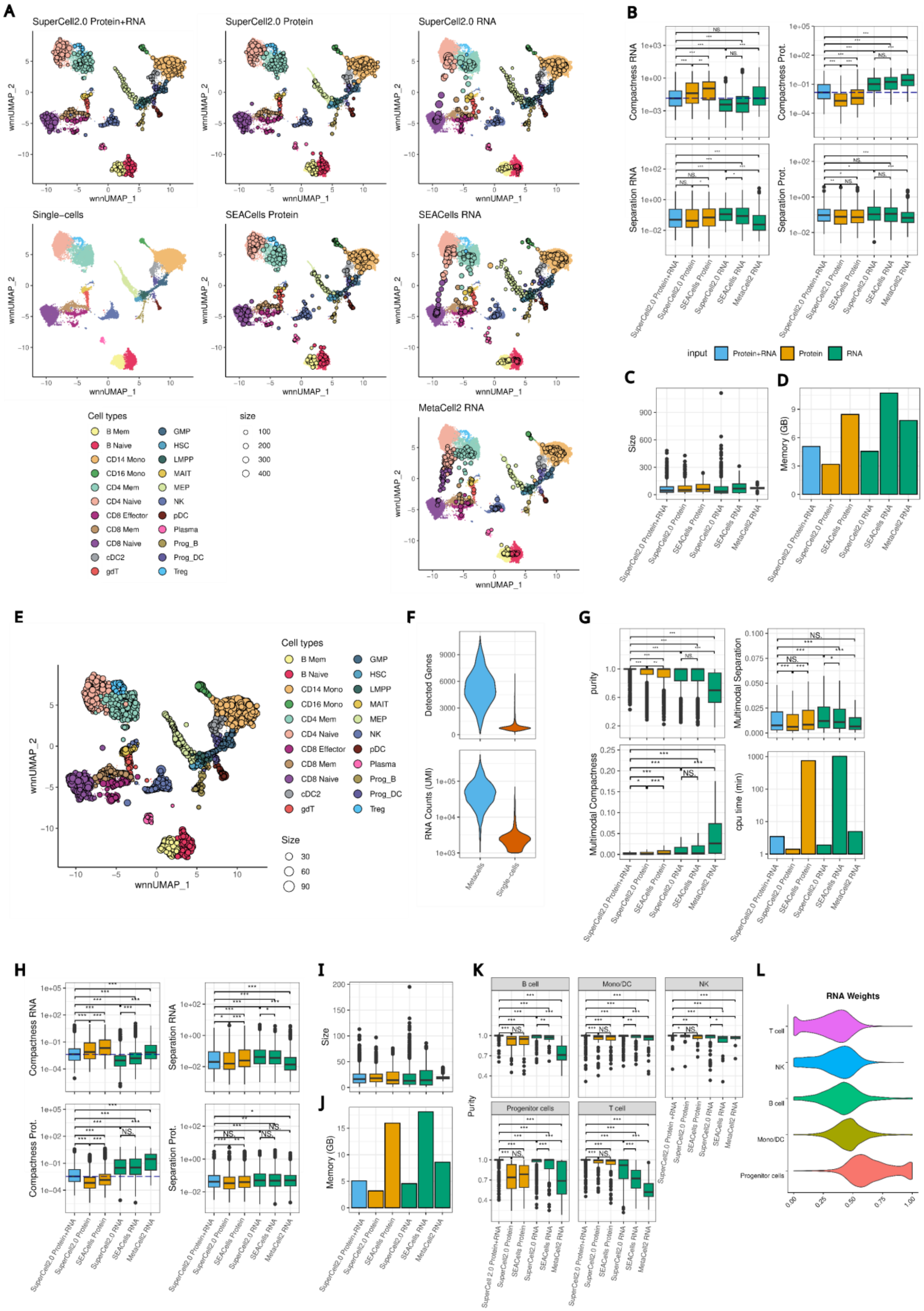


**Supplementary Figure 2: Benchmark of the unsupervised workflow of SuperCell2.0 on BM CITE-seq data. A** Coverage plots of the single-cell WNN UMAP of the metacells identified with

SuperCell2.0, SEACells and MetaCell2 using as input Protein with RNA, Protein only and RNA only (γ=75). **B** RNA and Protein compactnesses and separations (log scale) for metacells obtained with SuperCell2.0 (unsupervised), SEACells and MetaCell2 using as input Protein & RNA, Protein only and RNA only. The horizontal dashed lines in compactness boxplots correspond to the median of the metacell compactnesses obtained with SuperCell2.0 multimodal workflow. **C** Size distributions of the corresponding metacells. To note that MetaCell2 algorithm keeps metacell size between a range. **D** Memory peak usage of metacell identification for SuperCell2.0, SEACells and MetaCell2 using as input Protein & RNA, Protein only and RNA only. **E** Coverage plot of the single-cell WNN UMAP of metacells identified at γ=20 with the multimodal workflow of SuperCell2.0. **F** Detected gene and UMI count distributions of the metacells in **E** and single cells for the BM CITE-seq data. **G** Metacell purities, multimodal separations and compactnesses (outliers not shown), and CPU time of the tools SuperCell2.0, SEACells and MetaCell2 using as input Protein & RNA, Protein only and RNA only (γ=20). **H** RNA and Protein compactnesses and separations of the tools SuperCell2.0, SEACells and MetaCell2 using as input Protein & RNA, Protein only and RNA only (γ=20). The horizontal dashed lines in compactness boxplots correspond to the median of the metacell compactnesses obtained with SuperCell2.0 multimodal workflow. **I** Metacell sizes and **J** Memory peak usage of the tools SuperCell2.0, SEACells and MetaCell2 using as input Protein & RNA, Protein only and RNA only (γ=20). **K** Metacell purity in the major cell types of the BM-CITE-seq data. **L** Single-cell RNA weights resulting from the weighted nearest neighbors analysis of the single-cell BM CITE-seq data in each major cell type. Differences in metric distributions were tested using Wilcoxon tests In **B**, **G, H** and **K** with results grouped into (***) for p-values < 0.001, (**) for p-values < 0.01, (*) for p-values < 0.05 and no significant difference (NS.) for p-value > 0.05.

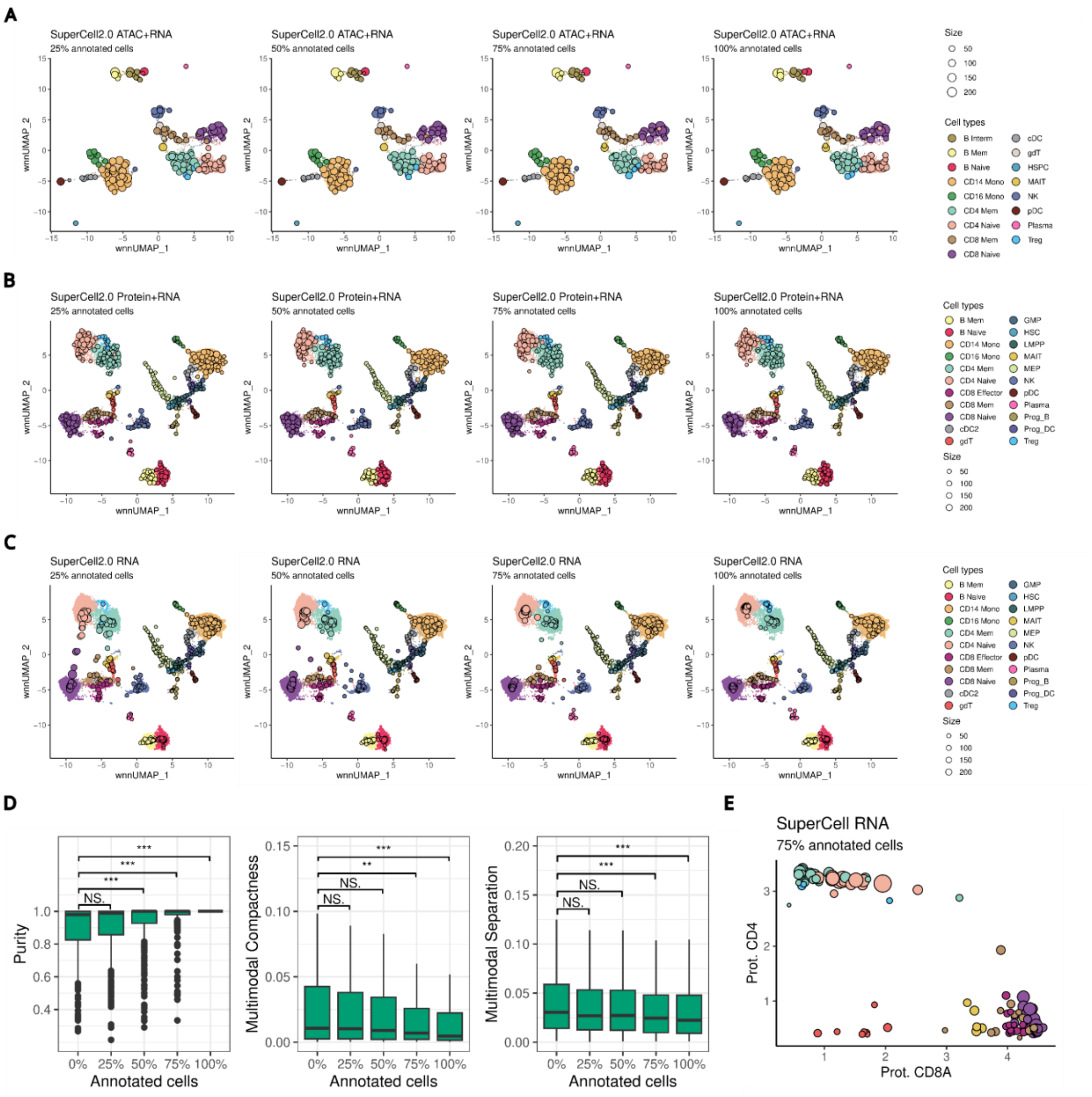


**Supplementary Figure 3: Benchmark of the semi-supervised workflow of SuperCell2.0. A** Coverage plot of single-cell WNN UMAP of PBMC 10x Multiome data of metacells identified with the semi-supervised workflow of SuperCell2.0 (ATAC+RNA, γ=75), using increasing percentage of annotated cells in input. **B** Coverage plot of single-cell WNN UMAP of BM CITE-seq data of metacells identified with the semi-supervised workflow of SuperCell2.0 (Protein+RNA, γ =75) using increasing percentage of annotated cells in input. **C** Coverage plot of single-cell WNN UMAP of BM CITE-seq data of metacells identified with the semi-supervised workflow of SuperCell2.0 using only RNA (γ=75) and increasing percentage of annotated cells in input. **D** Purity, multimodal compactness and separation of metacells identified at γ=75 and increasing percentage of annotated cells on the BM CITE-seq data with the supervised workflow of SuperCell2.0 using only RNA (γ=75). Differences in metric distributions with the unsupervised workflow were tested using Wilcoxon tests with results grouped into (***) for p-values < 0.001, (**) for p-values < 0.01, (*) for p-values < 0.05 and no significant difference (NS) for p-value > 0.05. **E** CD8A and CD4 protein expression in T lymphocyte metacells identified at γ=75 with SuperCell2.0 semi-supervised workflow using only RNA with 75% of annotated cells.

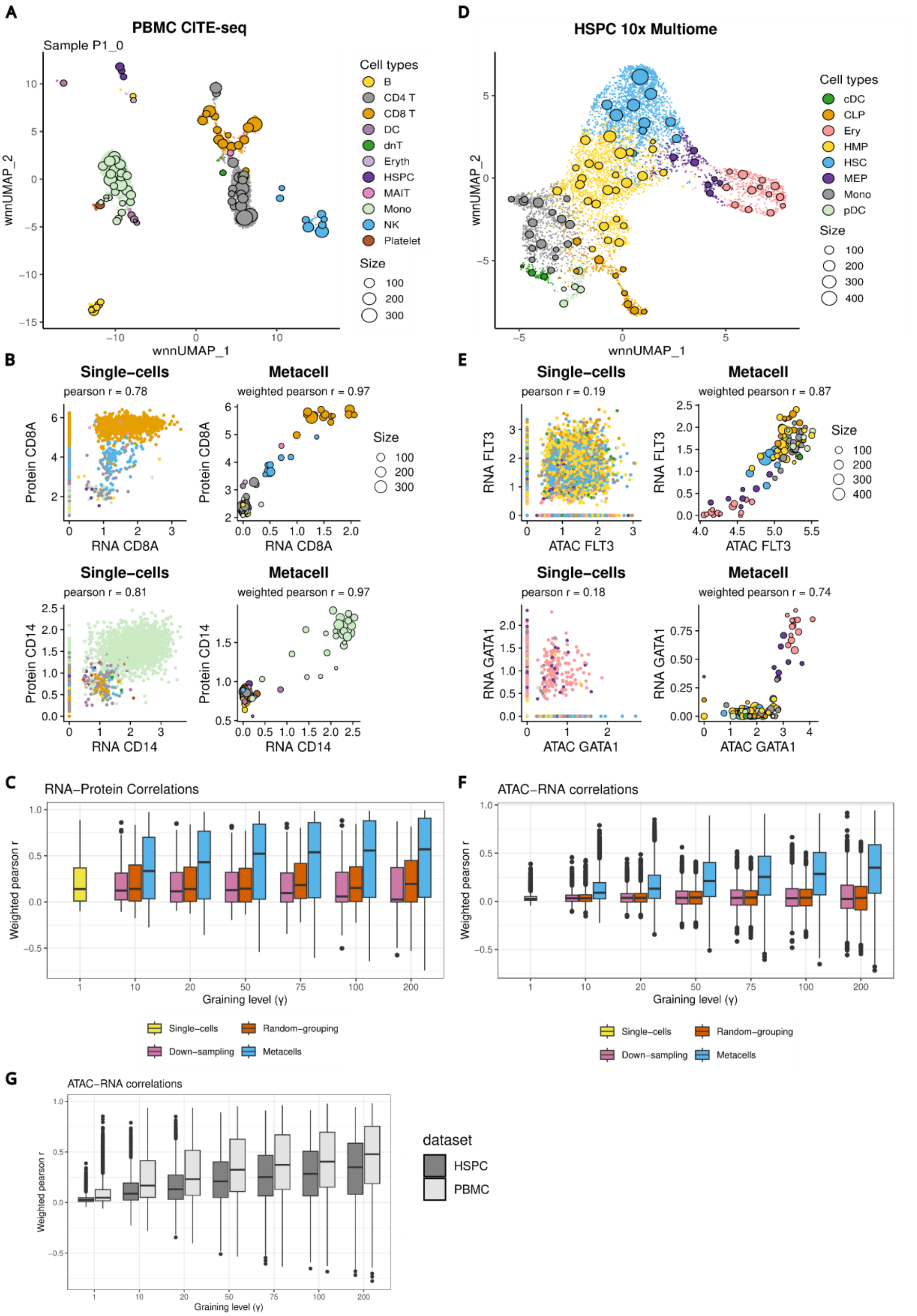


**Supplementary figure 4: Metacells improve intermodality consistency in PBMC CITE-seq and HSPC 10x Multiome datasets. A** Coverage plots of metacells identified by the unsupervised multimodal SuperCell2.0 workflow on the sample P1_0 of the human PBMC CITE-seq atlas at γ=75. **B** RNA and surface protein expression in single cells (left) and metacells (γ=75, right) for CD8A and CD14 in the BM CITE seq data. Same legend as in **A**. **C** Distribution of weighted Pearson RNA-Protein correlations for the 200 measured proteins in the PBMC CITE-seq data at increasing γ. Down-sampling (subsetting) and random grouping of single cells were performed as controls for each tested γ. **D** Coverage plots of metacells identified by the unsupervised multimodal SuperCell2.0 workflow on a human HSPC 10x Multiome dataset at γ=75. **E** Gene activity (ATAC) and RNA expression in single cells (left) and metacells (γ=75, right) for *FLT3* and *GATA1* in the HSPC 10x Multiome data. Same legend as in **D**. **F** Distribution of weighted Pearson gene activity - gene expression (ATAC-RNA) correlations at increasing γ for 4000 highly variable genes at the single-cell RNA level in the HSPC 10x Multiome data. Down-sampling (subsetting) and random grouping of single cells were performed as controls for each tested γ. **G** Gene activity (ATAC) and RNA correlation for 4000 highly variable genes at the single-cell RNA level at increasing γ in the HSPC and PBMC 10x Multiome datasets.

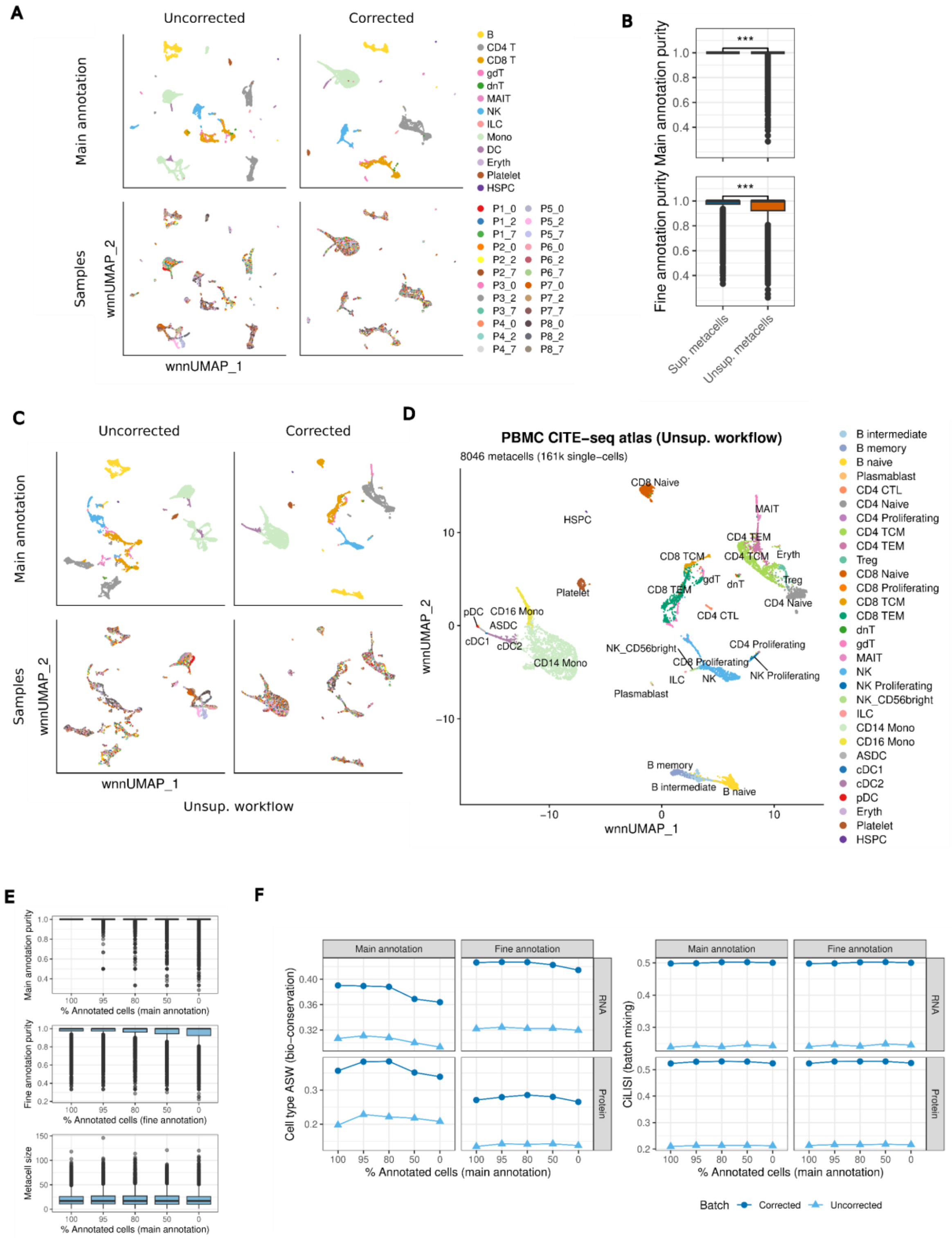


**Supplementary Figure 5: Semi-supervised metacells enable the integration of a PBMC CITE-seq atlas. A** WNN UMAPs of the integrated PBMC CITE-seq atlas before and after batch correction of RNA and protein latent space using the supervised workflow. Metacell are colored by main labels (aggregated single-cell labels, top panel) and samples (bottom panel). **B** Main and fine annotation labels purities of metacell obtained with the supervised and the unsupervised workflow. **C** WNN UMAPs of the integrated PBMC CITE-seq atlas before and after batch correction of RNA and protein latent space using the unsupervised workflow. Metacell are colored by main labels (aggregated single-cell labels, top panel) and samples (bottom panel). **D** WNN UMAP of the integrated PBMC CITE-seq metacell atlas using the unsupervised workflow. Metacells are colored by fine annotation. **E** Purity of main and fine annotation labels, as well as size of metacells obtained with decreasing percentage of annotated single cells. **F** Cell type ASW (cell type separation, left panel) and CiLISI (batch mixing, right panel) of integrated metacells using the semi-supervised workflow with increasing percentage of annotated cells. The metrics are computed using main and fine annotation labels in RNA and Protein latent spaces before and after batch correction.

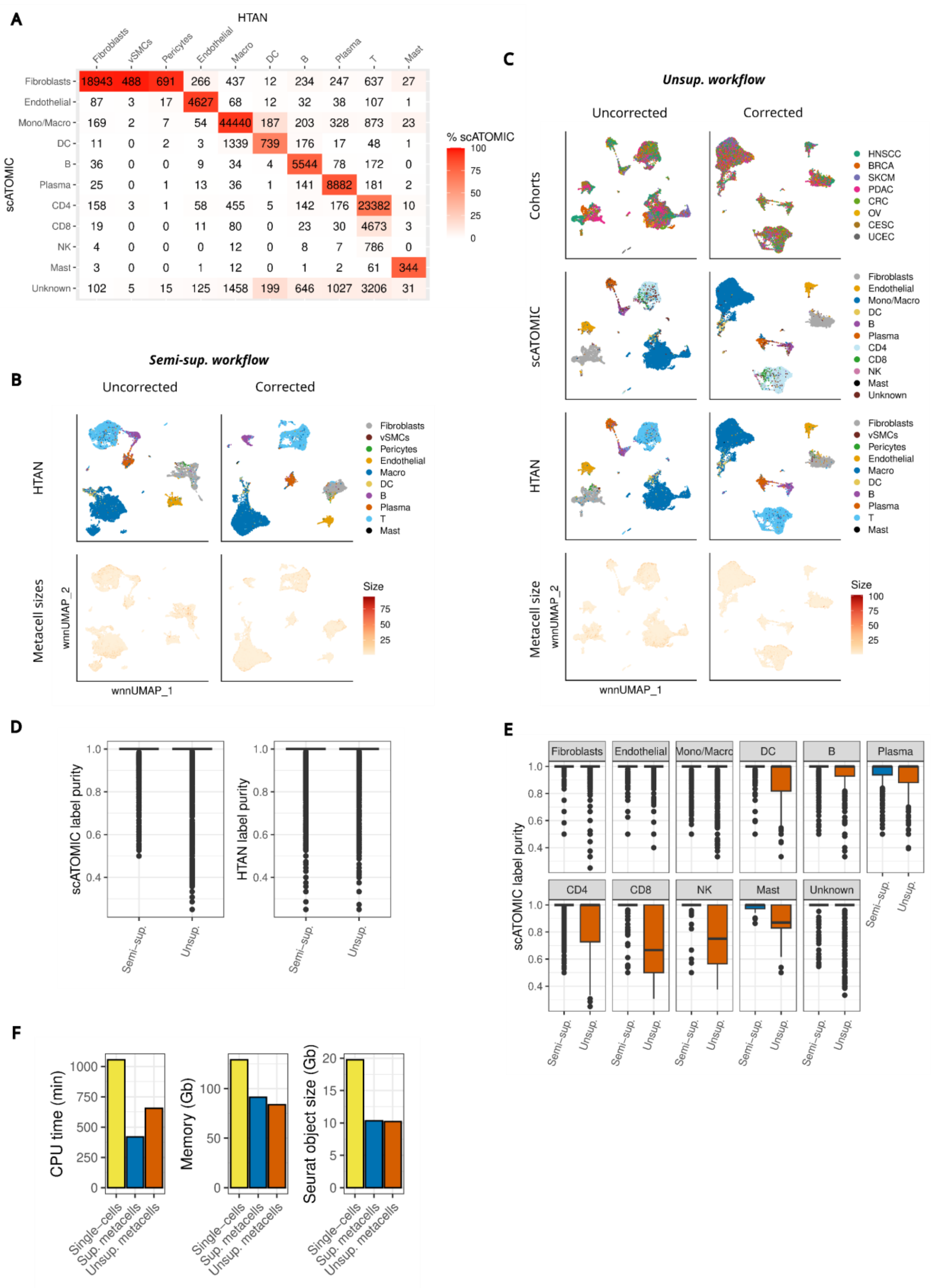


**Supplementary Figure 6: Semi-supervised metacells enable the integration of a TISME 10x Multiome atlas. A** Correspondence of single-cell annotations by scATOMIC with the original cell annotation of the HTAN data. **B** WNN UMAPs of the TISME 10x Multiome atlas before and after batch correction of RNA and ATAC latent space using the semi-supervised workflow.

Metacells are colored by original HTAN annotation (aggregated single-cell labels) and size. **C** WNN UMAPs of the TISME 10x Multiome atlas before and after batch correction of RNA and ATAC latent space using the unsupervised workflow. Metacells are colored by cancer type cohort, scATOMIC and original HTAN annotations (aggregated single-cell labels) and size. **D** Purity in scATOMIC and HTAN labels of metacell obtained with the semi-supervised and the unsupervised workflow. **E** Purity in scATOMIC label by cell type of the metacells obtained with the semi-supervised and the unsupervised workflow. **F** Computational load of the metacell semi-supervised workflow compared to an equivalent single-cell workflow.

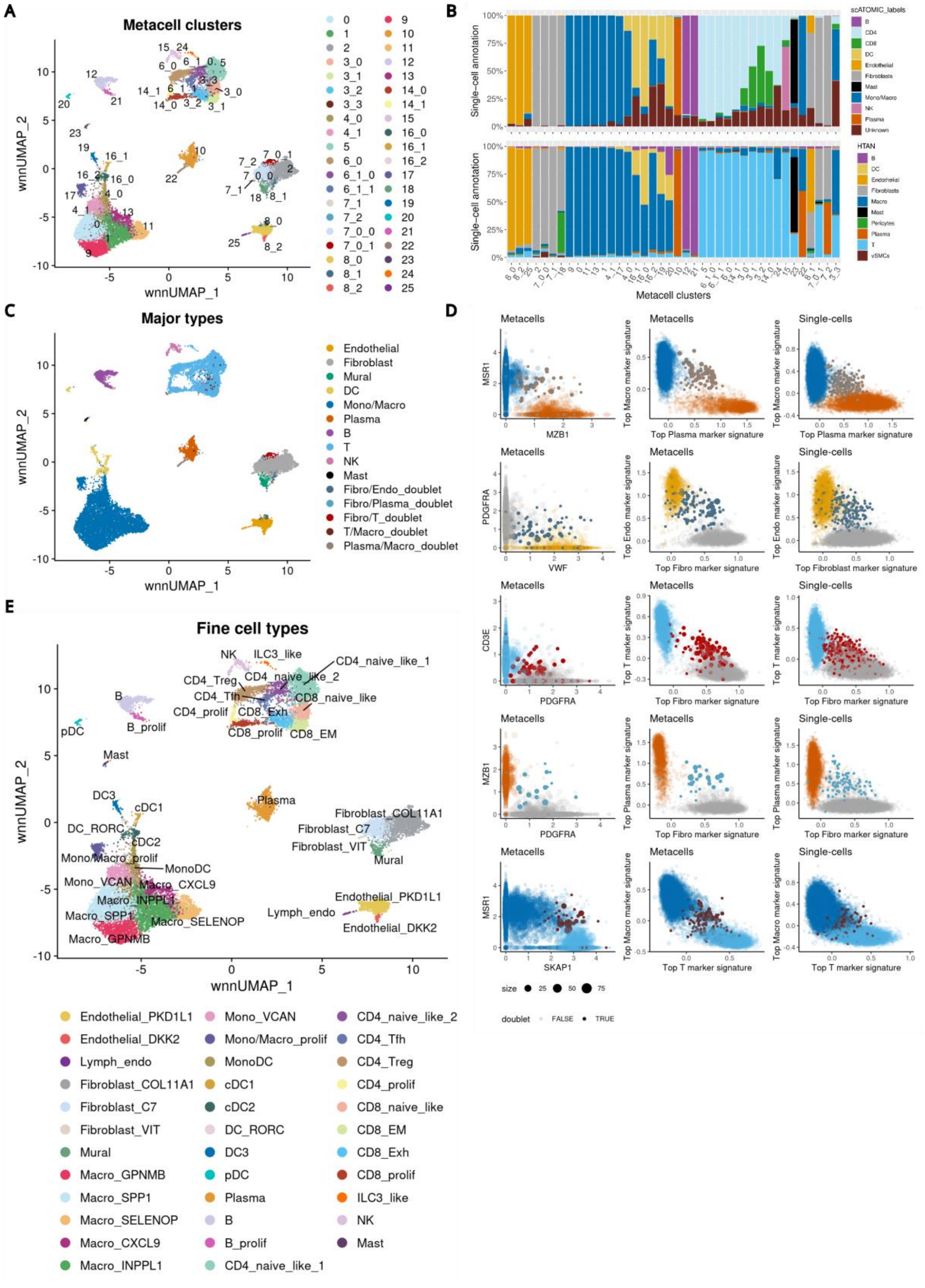


**Supplementary Figure 7: Annotation of the TISME Multiome atlas. A** WNN UMAPs of the TISME 10x Multiome atlas with metacells colored by multimodal clusters. Subclusters are denoted by an underscore ("_"). **B** Cell type composition of multimodal metacell clusters according to single-cell HTAN and scATOMIC annotations. **C** WNN UMAPs of the TISME 10x

Multiome atlas with metacells colored by major cell type annotation. **D** Scatter plots of marker gene expression (metacell) and signature scores (metacells and single-cells) of major types used to identify clusters of doublets. Metacells and single cells are colored by major cell type annotation as in **C**. **E** WNN UMAPs of the TISME 10x Multiome atlas with metacells colored by fine cell type annotation (metacell clusters of doublets are discarded).

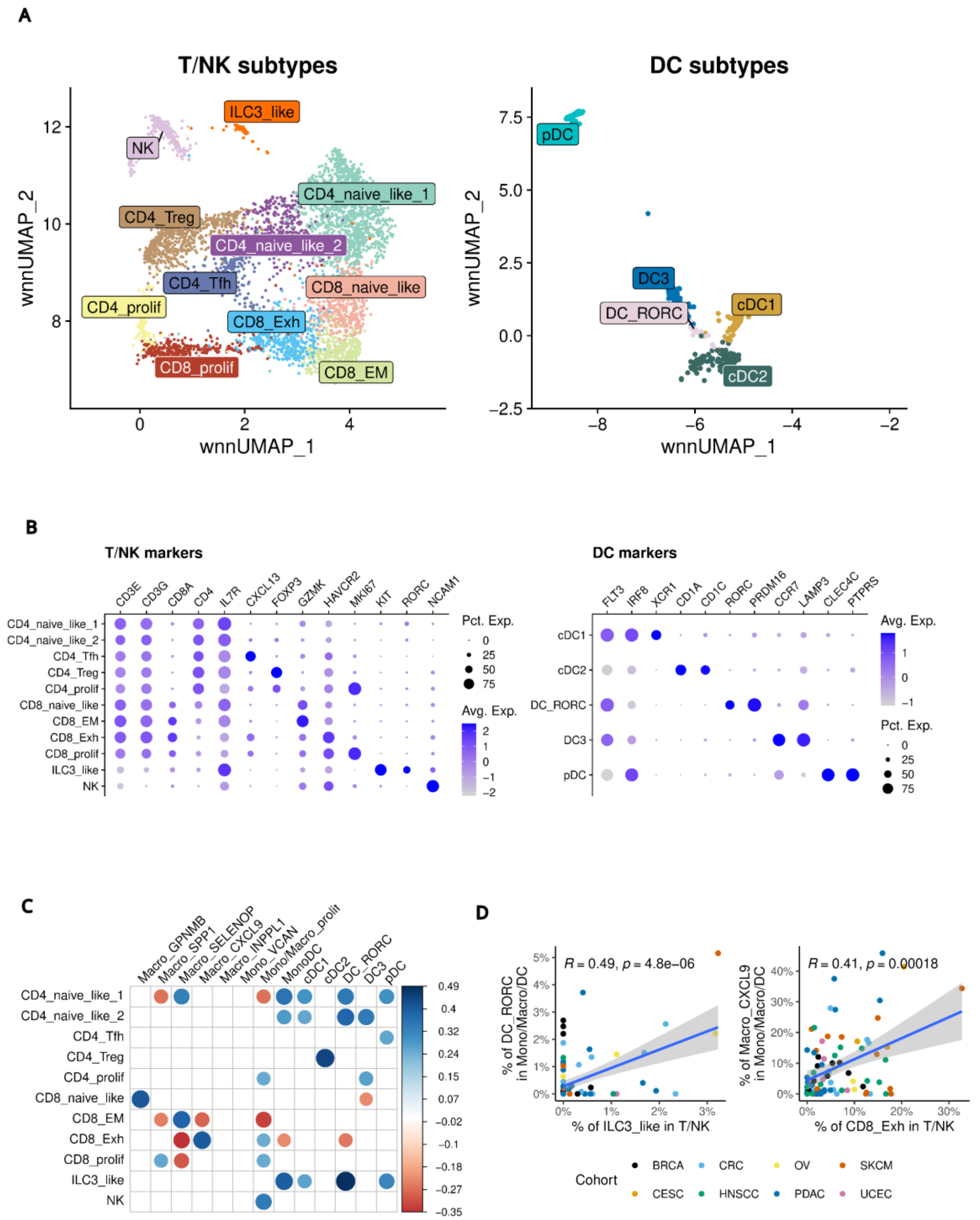


**Supplementary Figure 8: Analysis of the TISME atlas. A** WNN UMAPs of the T/NK (left panel) and DC (right panel) subtypes. **B** Dot plots of selected RNA markers for the T/NK (left panel)

and DC (right panel) subtypes. **C** Significant correlation of abundances of T/NK with Mono/Macro/DC subtypes (p-value of Pearson correlation test < 0.05). **D** Selected significant correlations of abundances of T/NK with Mono/Macro/DC subtypes.

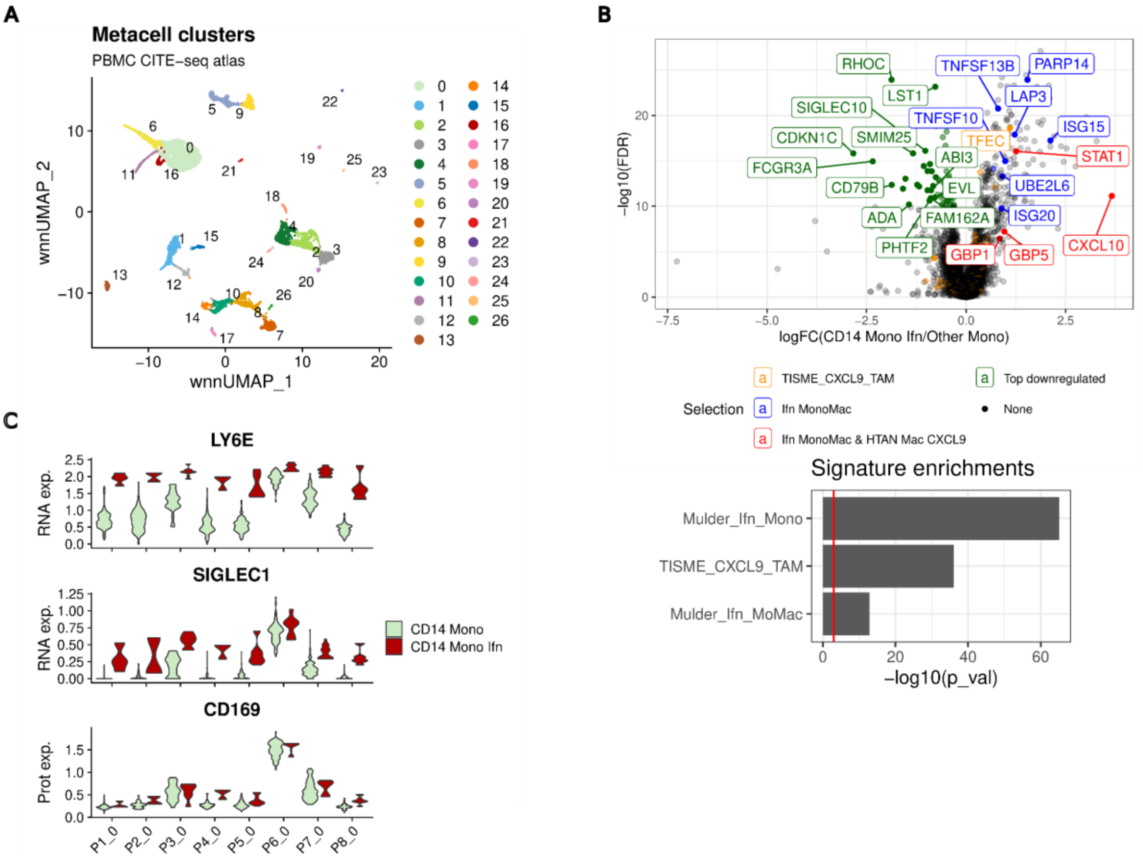


**Supplementary Figure 9: Metacell analysis of interferon-primed CD14 monocytes in blood from healthy donors. A** WNN UMAP of the PBMC CITE-seq metacell atlas. Metacell are colored by clusters. **B** Upper panel**:** Volcano plot of DEG between interferon-primed monocytes and other monocytes. Top genes of signatures are highlighted. Lower panel: gene set enrichments of interferon-primed CD14 monocytes markers for selected signatures from the literature (hypergeometric test). **C** Violin plot of *LY6E*, *SIGLEC1* RNA expression and CD169 (encoded by *SIGLEC1*) protein expression in metacells for each healthy donors (day 0 of the vaccination trial) in CD14 monocytes compared to interferon-primed CD14 monocytes clusters.

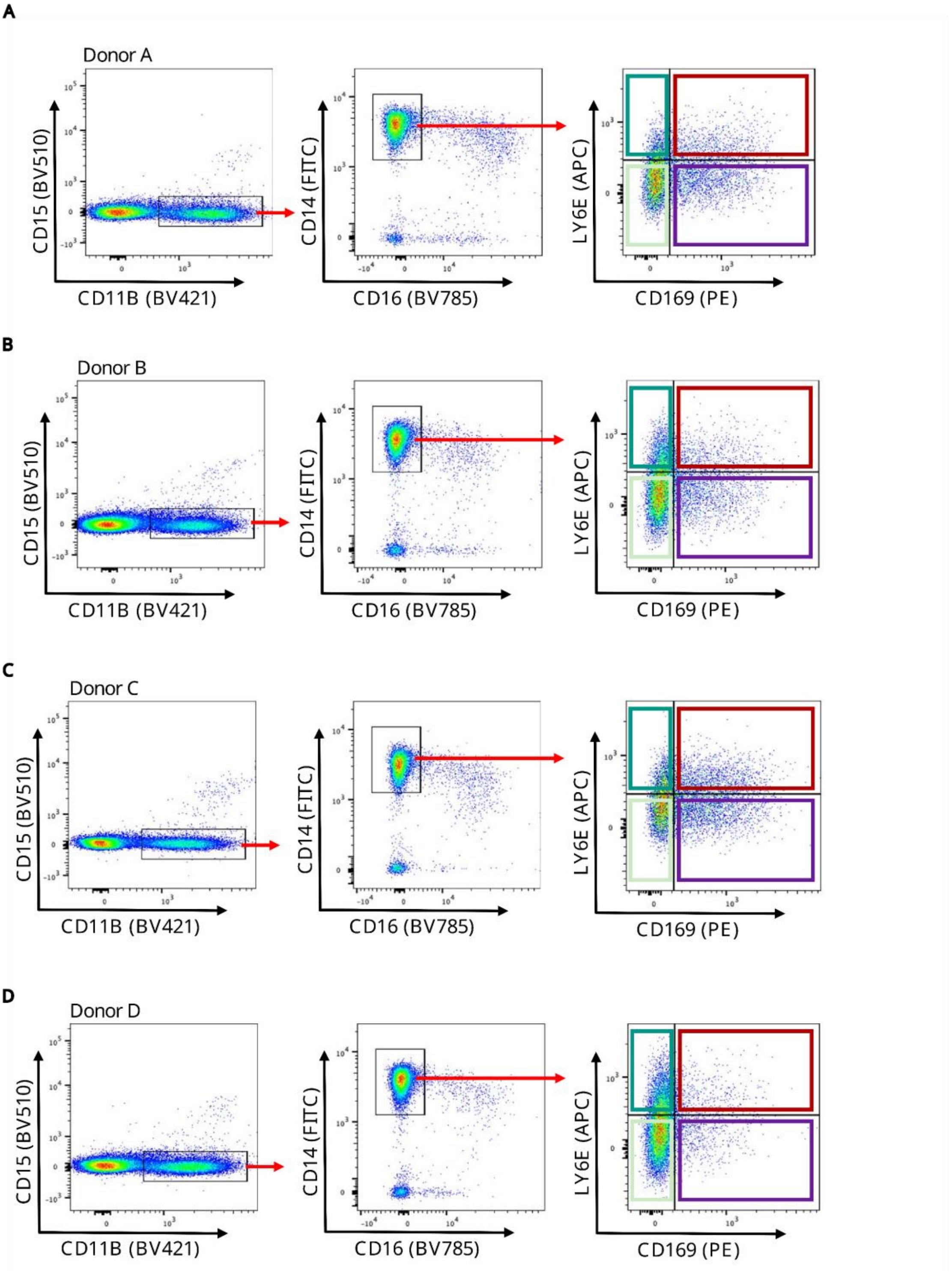


**Supplementary Figure 10: FACS analysis of blood monocytes from four healthy donors.** FACS gating strategy and corresponding scatter plots of cytometry data to sort and analyze CD14 monocytes subtypes from the healthy blood donors **A**, **B**, **C** and **D**
